# Hybrid supramolecular-covalent bioresin promotes cell migration and self-assembly in light-based volumetric bioprinted constructs

**DOI:** 10.1101/2025.01.06.631505

**Authors:** Marc Falandt, Paulina Nunez Bernal, Alessia Longoni, Maj-Britt Buchholz, Pere Català Quilis, Klement Widmann, Mario Barrera-Roman, Jos Malda, Tina Vermonden, Anne Rios, Riccardo Levato

## Abstract

There is an increasing need for novel biomaterials compatible with advanced biofabrication technologies, which also permit cells to remodel their microenvironment. This remodelling is crucial for maturing tissue constructs into fully functional tissue replacements. Recent progress in supramolecular chemistries has allowed for the production of dynamic biomaterials. Their properties enable bonds to be reversibly broken by cells, facilitating processes requiring morphological changes or migration, crucial for tissue development and homeostasis. Here, we present a one-of-its-kind gelatin-based hybrid covalent/supramolecular biomaterial. We demonstrate the advantage of adding supramolecular-reactive moieties on covalent materials, over covalent bonds alone, in facilitating processes such as cell growth, migration, spreading and organoid proliferation. This is exemplified by enhanced MSC and T cell migration and improved vascular network formation in hybrid hydrogels over covalent-only materials. The combination of supramolecular and covalent bonds further enabled control over photocrosslinking, allowing the use of the material in volumetric bioprinting of complex structures with high shape fidelity. As a proof-of-concept we bioprinted complex breast-like structures from encapsulated normal breast cell lines with a tumor organoid core. We demonstrated that engineered T cells can migrate large distances into the breast tissue, specifically targeting tumor cells.

**Graphical abstract:** 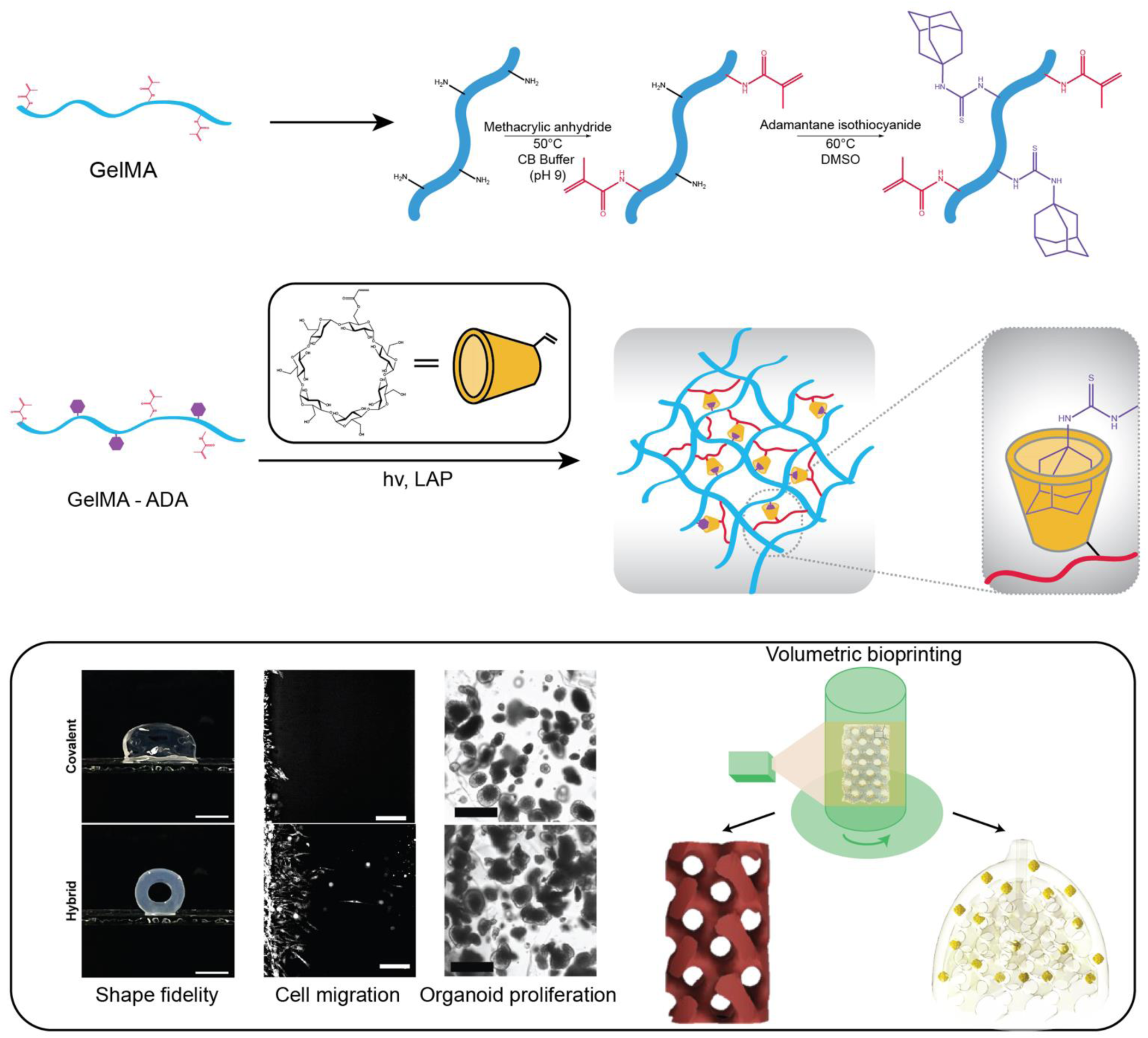

## Introduction

The extracellular matrix (ECM) is amenable to dynamic changes triggered by cellular spreading, re-organization, growth, differentiation and migration, fundamental for functional tissue development, morphogenesis and homeostasis^1–3^. Reciprocally, the ECM composition, spatial arrangement and mechanics influence cell behavior and fate^2–5^. To generate functional in vitro 3D tissue models, it is therefore important that hydrogels used for tissue engineering and cell culture are able to recapitulate the dynamic nature of the ECM. While biofabrication methods provide the ability to generate complex physiological architectures^6–8^, many naturally-derived biomaterials that enable cell-induced remodelling, often do not possess the required shape fidelity and stability post-printing^2,9–13^.

Many strategies aim to increase a biomaterials permissiveness to native cellular processes, through stimuli-responsiveness or including hydrolytic or cell-mediated enzymatic degradation. However, those processes may lead to a loss of stability and shape fidelity over time. Moreover, they are often one-directional in nature and therefore not able to capture the dynamic characteristics of the native extracellular matrix.

Much research has, therefore, been executed on the development of biomaterials for biofabrication that possess both, desirable printability and long-term shape fidelity, as well as permissiveness for cellular processes requiring cellular growth, morphological changes or migration^14–18^. This has shifted the focus from *static* material properties to *dynamic* environments^5,18^. Reversible chemistries have herein played an important role to create materials with fast relaxation kinetics such as supramolecular materials. Those employ non-covalent bonds that can be reversibly broken by cells to migrate and expand in the material and quickly thereafter re-form, ensuring a maintained shape fidelity^2^. Such supramolecular materials have, for example, shown great success in facilitating vascular assembly and invasion or for drug delivery^14,19,20^.

Recently, supramolecular hydrogels have furthermore been used in the context of organoid culturing demonstrating the influence of dynamic culture environments on promoting kidney organoid glomerulogenesis^21^, intestinal organoid morphogenesis^22^ and trophoblast organoid differentiation^23^. Supramolecular materials might therefore be a promising approach to replace Matrigel. This gold-standard for organoid culture is a biochemically poorly defined matrix^21,24,25^ with high batch-to-batch variability, which can hardly be modified to accommodate the requirements of different cellular processes or cell types^26^.

However, purely supramolecular materials might not provide the desired mechanical strength long-term stability and supramolecular bond formation cannot be spatially controlled. That is why dual/secondary crosslinking mechanisms have been proposed to tune mechanical strength and enable light-induced spatial selective crosslinking for bioprinting^27,28^.

Combining the advantageous dynamic nature of supramolecular interactions with the highly controllable light-induced covalent crosslinking of methacrylated groups, we here present for the first time the synthesis and application of a covalent/supramolecular hybrid gelMA for the use in tissue engineering and biofabrication.

We characterize, the beneficial effects of supramolecular bonds as additional feature on a gelatin covalent backbone, on diverse biological processes such as cell migration and organization, vascular network formation and organoid growth. These combination with the photocontrollable covalent bonds thereby enabled the use this material for light-based volumetric bioprinting. This yielded prints of complex shapes and intricate hollow features at high shape fidelity and stability. Finally, we utilize our hybrid material for generating a proof-of-concept co-culture model of the human breast containing normal epithelial and cancerous cells. We demonstrate its use for studying tumor specific targeted migration of engineered T cells.

## Results

### Hybrid supramolecular hydrogels form reversible host-guest interactions between acrylated β-cyclodextrin and adamantane

Gelatin methacryloyl (gelMA), was used a platform for further modification. This material is considered a golden standard in various bioprinting modalities given its mechanical versatility and ease of production at different degrees of methacrylic modification^11–13,29^. This collagen-based biomaterial also provides an extracellular matrix (ECM)-like environment, rich in cell adhesive motifs. It has therefore been used to culture a wide range of biological units (i.e. cells, aggregates and organoids) in the fields of tissue engineering and biofabrication^11,30–35^.

Here, we synthesized gelMA with different degrees of methacrylation (DoM) to yield hydrogels with various network densities and thus, mechanical properties. The remaining lysines on the gelatin backbone were then functionalized with adamantane isothiocyanate (AITC) until a 100% degree of functionalization (DoF) of all lysine residues was reached. This provided us with a library of covalent and hybrid gelMA polymers with a DoMs of 40, 60, 80, and 100%. These photosensitive materials could be photopolymerized in the presence of 0.1 w/v% lithium phenyl(2,4,6-trimethylbenzoyl)phosphinate (LAP).

In order to study the formation of hydrophobic supramolecular interactions of the AITC-functional groups in combination with the acrylated β-cyclodextrin (Ac-β-CD), we assessed hydrophobicity differences between all material formulations with and without addition of Ac-β-CD. For that we performed a Nile-Red assay, which indicates increases in hydrophobicity through increased fluorescence. We observed, that the hybrid material, prior to the addition of Ac-β-CD, was, as expected due to the hydrophobicicyt of AITC-groups, more hydrophobic than covalent gelMA. However, when Ac-β-CD was added, the hybrid material showed a decrease in hydrophobicity. Conversely, covalent gelMA showed an increase of hydrophobicity upon the addition of the Ac-β-CD. This indicates, that the hybrid material forms host-guest interactions between hydrophobic adamantane and the hydrophobic pocket of the Ac-β-CD, creating a more hydrophilic material (**Extended Fig 1**). Hence, the AITC-functionalization and addition of Ac-β-CD led to the spontaneous formation of both a covalent methacrylate-methacrylate (MA-MA) network and supramolecular host-guest interactions between adamantane and Ac-β-CD (**Extended Fig 2a**).

**Figure 1.**
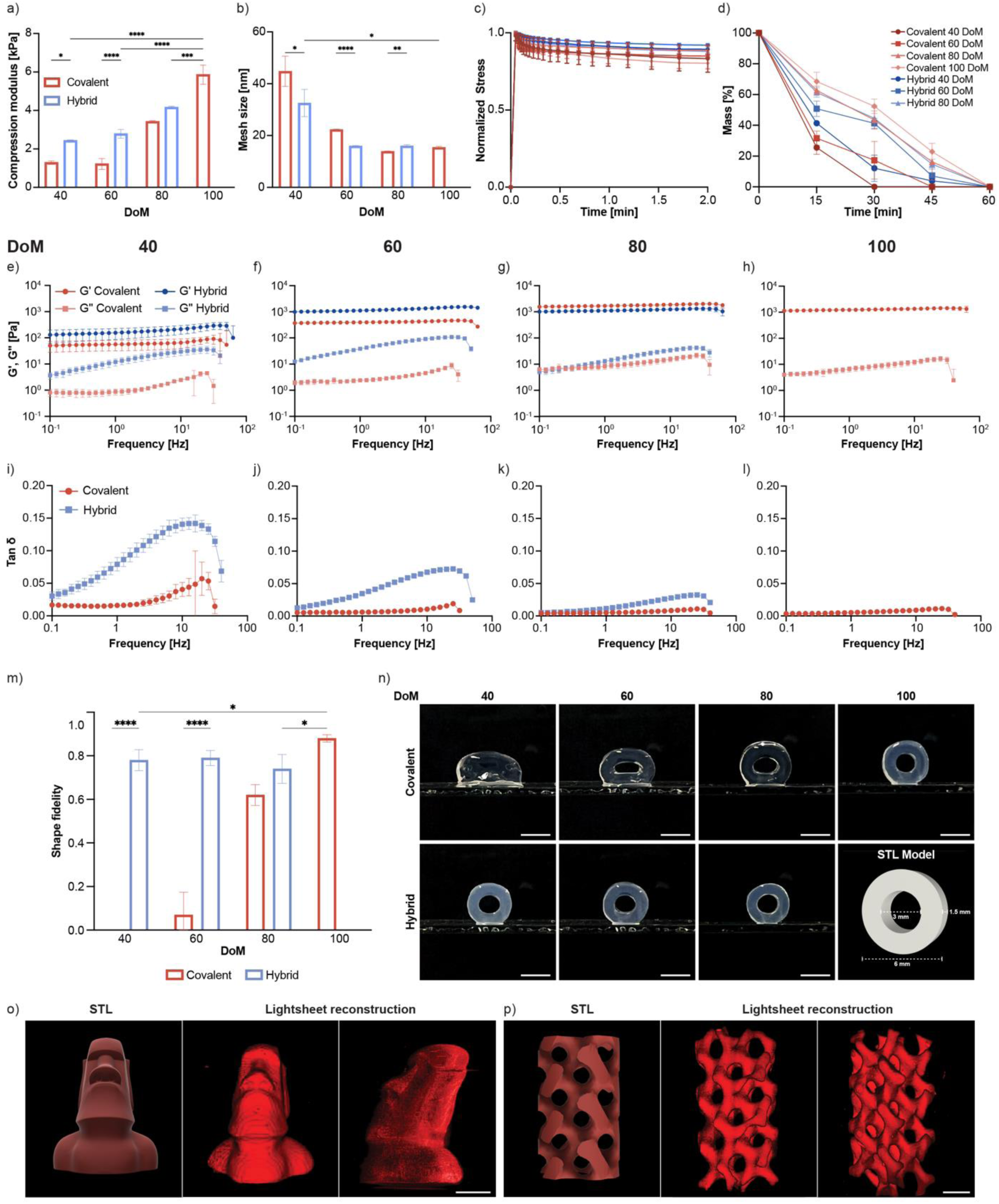
Mechanical and physical characterization of 5 w/v% covalent and hybrid bioresins with varying DoM (40, 60, 80, and 100) for volumetric printing. a) Compressive modulus of crosslinked hydrogels (n = 3 to 8). b) Calculated mesh size in nm derived from the max G’ of the frequency sweep for different hydrogel formulations (n = 3). c) Normalized stress-relaxation evolution graphs (n = 3). d) Enzymatic material degradation in collagenase solution at 37°C (n = 3). e, f, g, h) Rheological frequency sweep (0.1 to 100 Hz) of different hydrogel formulations after photocrosslinking (40 DoM, 60 DoM, 80 DoM, 100 DoM respectively) (n = 3). i, j, k, l) Tan delta as calculated from the frequency sweep of the independent hydrogel materials (n = 3). m) Normalized shape fidelity of circular volumetric printed models using the different hydrogel formulations (n = 3). n) Volumetrically printed models for shape fidelity calculations with the intended STL model as comparison, scalebar = 5 mm. o, p) CAD model and lightsheet reconstruction of the Hybrid 60 DoM hydrogel, showing an Eastern Island statue and a gyroid model respectively, scalebar = 2 mm.

**Figure 2.**
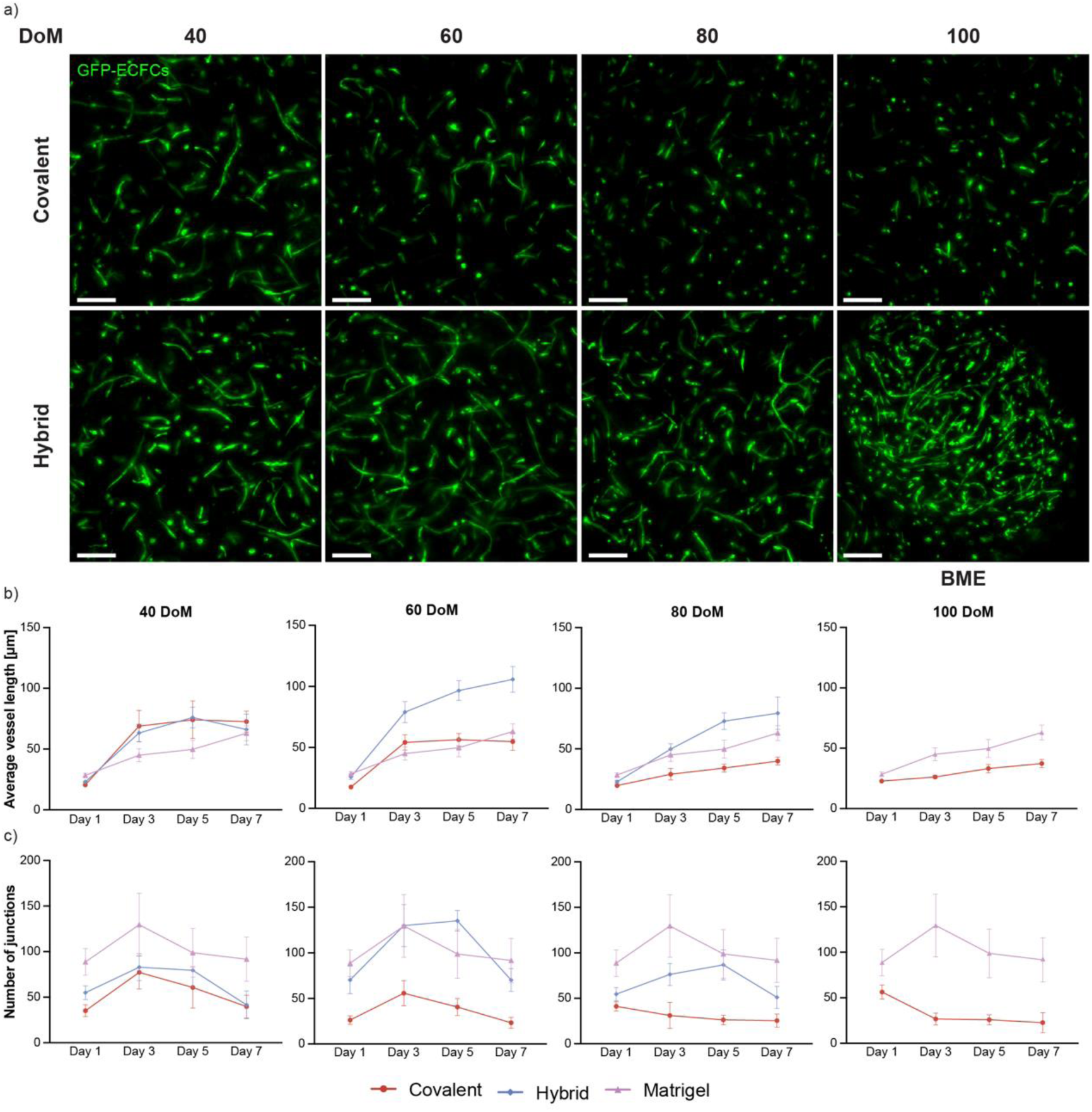
Vascular network formation within hybrid bioresins. a) Fluorescent images showing ECFCs (green) embedded in covalent and hybrid (DoM: 40, 60, 80 and 100) hydrogels after 7 days in 3D culture with hMSCs (1:3 ratio of ECFCs to hMSCs), with BME droplets as a control, golden standard material. Scale bar = 200 µm. b) Average vessel length and c) total number of junctions of interconnected ECFCs, each graph showing a different DoM hydrogel with both covalent and hybrid counterparts, as well as BME.

### Hybrid supramolecular hydrogels display viscoelastic properties

Our library of covalent and hybrid gelMA formulations provides materials with a highly tunable range of mechanical properties, largely dependent on the different DoM of the materials. They further exhibit interesting viscoelastic properties depending on the degree of AITC functionalization.

As expected, a significant increase of the compressive modulus was observed in all hydrogel formulations with increasing DoMs. This suggests that the bulk stiffness of both covalent and hybrid gelMA hydrogels is largely determined by the covalent gel network (**Fig. 1A1a**). Interestingly, a significant increase in the compressive modulus of hybrid materials compared to the covalent gelMA controls with the same DoM could be observed. Here, the additional supramolecular interactions also appear to contribute to the bulk stiffness of the hydrogel by creating a more densely crosslinked network compared to the covalent counterparts (**Figure 1B1b**). The spontaneous formation of these supramolecular interactions and their contribution to material stiffness was confirmed, when the hybrid material was crosslinked without the addition of Ac-β-CD. This resulted in a less pronounced the increase of the compression modulus (**Extended Fig. 2b**). Similarly, covalent-only gels (without adamantane functionalization), displayed less pronounced changes in Young’s modulus upon the addition of Ac-β-CD (**Extended Fig. 2c**). This suggests, that it is the host-guest interactions present in the hybrid material that provide the additional mechanical strength. Those results correlate with the measured mesh sizes, which show an increased hydrogel network density in hybrid formulations in presence of Ac-β-CD (**Extended Fig. 2D,E2d,e**). The increased network density is further demonstrated by the significantly lower swelling ratios of the hybrid 40 and 60 DoM formulations compared to their covalent counterparts (**Extended Fig. 2f**). Interestingly, 80 DoM hybrid gel shows strong similarities to the covalent only formulation. This is likely due to the lower supramolecular interaction ratio. Further, the fact that the addition of Ac-β-CD creates a larger mesh size compared to those created by the covalent-only interactions at higher DoMs could contribute to these similarities.

Despite the bulk stiffness differences, the elastic behavior and the slow stress-relaxation behavior of the hybrid material did not significantly differ to that of the covalent gelMA formulations (**Fig. 1C1c, Extended Fig. 2g,i-l**). The same holds true for the creep characteristics, suggesting a similar bulk mechanical behavior (**Extended Fig.2m-p**).

To show the incorporation of the Ac-β-CD with the adamantane into the covalent methacrylate network, a sol-fraction assay was performed (**Extended Fig. 2h**). This showed no significant differences in any of the conditions except 80 DoM, which indicates, that Ac-β-CD is properly crosslinked within the network without a loss of components. Further, enzymatic degradation of the hydrogel formulations showed no loss of biodegradability in hybrid formulations (**Fig. 1d**). However, slower degradation patterns were observed in hybrid materials as opposed to covalent controls. Both, the covalent gelMA with lower DoMs (40 and 60), showed complete degradation within 45 minutes. All hybrid formulations and high DoM covalent gels degraded only within 60 minutes, confirming that the presence of supramolecular interactions provides for a higher structural integrity of the hybrid hydrogel. As expected, the stiffer gels degraded slower, implying better long-term stability for cell cultures, whereas most soft materials do not provide sufficient structural integrity throughout long-term cell culture.

However, the aim was to produce hydrogels that not only provide long-term stability and shape fidelity but were also malleable by cells. We consequently sought out to better understand the reversible dynamic environment provided by the supramolecular bonds and through their viscoelastic behavior. Therefore, a frequency sweep within the viscoelastic region, as determined via amplitude sweep (**Extended Fig. 2q-t**), was performed. Hybrid materials exhibited higher storage moduli (G’) and loss moduli (G’’) than the covalent controls (**Fig. 1e-h**). Particularly, increased supramolecular substitution led to a significant increase in the loss modulus. This resulted in an increase in the Tan δ of the material (**Fig. 1i-l**) in line with previously described supramolecular networks composed of hyaluronic acid and PEGDA-modified fibrinogen^36^. Hence, supramolecular hybrid materials provide a stiffer environment, more suitable and stable for long-term cell culture as compared to covalent gels. As shown by the increased loss modulus at frequences relevant for forces exerted by cells during migration and adhesion, this material displays viscous behavior during these processes. Taken together, the environment provided by our hybrid material is permissive to cellular forces, all the while exhibiting no significant differences in photocrosslinking kinetics or amplitude sweep (**Extended Fig. 2u-x**).

### Hybrid biomaterials display enhanced shape fidelity upon light-based bioprinting

With the improved mechanical stability of our hybrid materials, shape fidelity was assessed using volumetric bioprinting. Hollow tubes of 3 mm inner diameter and 1.5 mm wall thickness were printed from all bioresin formulations and shape fidelity was assessed as measure of height versus width of the opening. As expected, hybrid materials exhibited a pronounced and significantly improved shape-fidelity compared to the covalent gelMA controls. Even for materials with low DoM (40 or 60 DoM), which normally do not provide shape-retaining printed constructs, the substitution of supramolecular bonds was sufficient to create self-supporting structures. Surprisingly, despite the higher compressive modulus of covalent 80 DoM hydrogel, both the hybrid 40 and 60 DoM displayed a better shape fidelity (**Fig. 1m,b**). Moreover, we were able to print more complex architecture featuring more intricate positive structure **(Fig. 1o)** and gyroidal negative channel networks **(Fig. 1p)** from hybrid materials as previously published with gelMA^34,35,37^. We observed an improvement in resolution for hybrid materials compared to the covalent gels. This is most pronounced for printing of positive features. Stiffer gels exhibited a better resolution than the softer materials, while no significant differences could be observed for printing of negative features (**Extended Fig. 3**).

**Figure 3.**
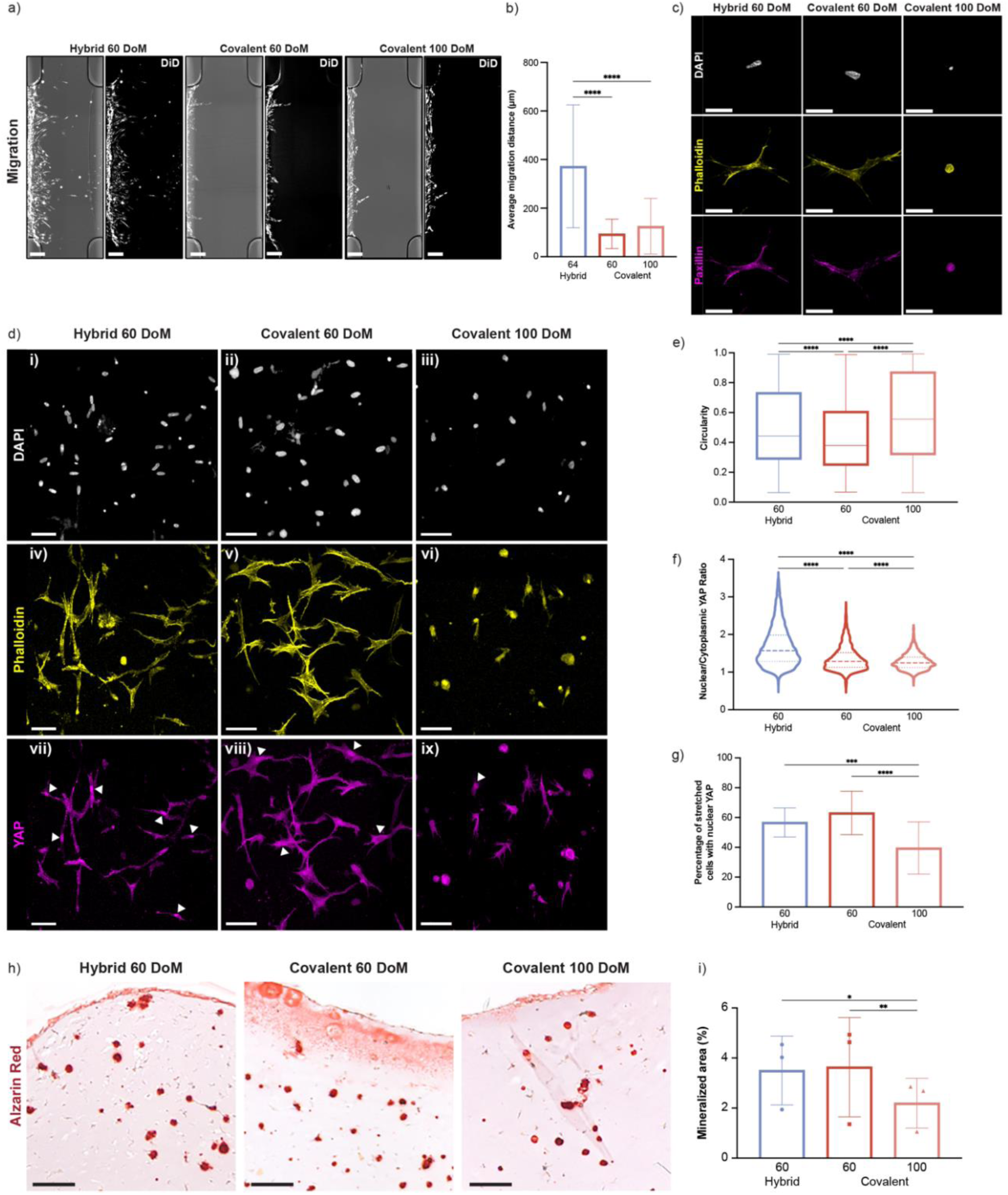
Mechanobiological cellular responses within hybrid bioresins. a) hMSC migration assay through hybrid 60 DoM and covalent 60 and 100 DoM hydrogels towards a chemoattractant (bFGF) chamber after 3 days in culture (left panel shows merged brightfield and DiD-labelled cells, right panel shows DiD-labelled cells only, scale bars = 200 mm). b) Quantification of hMSC migration distance. c) Representative confocal images of DAPI (grey), F-actin (yellow) and paxillin (magenta). Scale bar = 25 µm. d) Representative confocal images of DAPI (i-iii), phalloidin (iv-vi) and YAP (vii-ix) expression in hMSCs cultured in hybrid 60 DoM (i,iv,vii) and covalent 60 (ii,v,viii) and 100 (iii, vi,ix) DoM hydrogels after 3 days (scale bars = 50 µL). e) Quantification of cell circularity. f) YAP expression quantification expressed as the ratio of nuclear to cytoplasmic YAP localization. g) Quantification of stretched cells (circularity < 0.5) with nuclear translocation (Nuc/Cyto YAP ratio > 1.25). h) Representative images Alizarin Red staining of MSCs cultured in hybrid 60 DoM and covalent 60 and 100 DoM and hydrogels after 21 days of osteogenic differentiation. i) Corresponding quantification of Alizarin Red positive staining of calcified matrix.

### Hybrid gelMA formulations outperform covalent gels in vascular network formation

To ensure proper oxygen and nutrient, as well as waste transport, most tissues in our bodies are highly vascularized reaching every last corner of our body through different scales of vessels cumulating in 10 µm scale microcapillaries^38^. This ensures, that metabolic needs are met throughout the tissue volume and is therefore essential to take into account in tissue engineering^38–41^. Despite advancements in fabrication technology however, the generation of fully vascularized tissue engineered constructs, particularly microcapillaries still poses a significant challenge^42–44^. Strategies aiming to induce neo angiogenesis in vitro include the incorporation of cell instructive cues and mechanical stimulation through flow^45–48^. However, key to the success of such strategies lies in the permissiveness of the used biomaterial to cell infiltration, migration and vessel formation.

We, therefore, tested whether endothelial cells would be able to generate vascular networks in our hydrogel formulations. Fluorescent endothelial colony forming cells (ECFC) were embedded in covalent and hybrid gels, as well as BME in co-culture with human mesenchymal stem cells at a ratio of 1:3. Within a week of culture, ECFCs formed intricate vascular networks in 40 and 60 DoM covalent gels, BME and all hybrid formulations (**Fig. 2a, Extended Fig. 4**). As expected, due to the decreased mesh size with increasing DoM, average vessel length and number of junctions decreased with increasing DoM for covalent gels. Vessel length and number of junctions did not differ significantly between covalent and hybrid gels of 40 DoM, likely attributed to its already big mesh sizes. For higher DoM gels, however, we observed an increase of average vessel length and number of junctions of the hybrid gels compared to covalent formulations (**Fig. 2b,c**). This indicates that hybrid gels might be more permissive to cellular self-assembly leading to vascular network formation than covalent materials. Interestingly, highest vessel lengths and number of junctions were not observed in the softest of the hybrid gel formulations but rather in the slightly stiffer 60 DoM formulation. Similar observations have recently been reported by Wei et al^14^ in dynamic supramolecular collagen-hyaluronic acid hydrogels. They observed endothelial cells forming the largest tubular lumens and longest vessel distances in supramolecular hydrogels of medium plasticity rather than high plasticity. They attributed this to a better balance between cell adhesion and contraction in hydrogels displaying moderate plasticity, which could explain the improved vascular network formation in 60 DoM over 40 DoM formulations.

**Figure 4.**
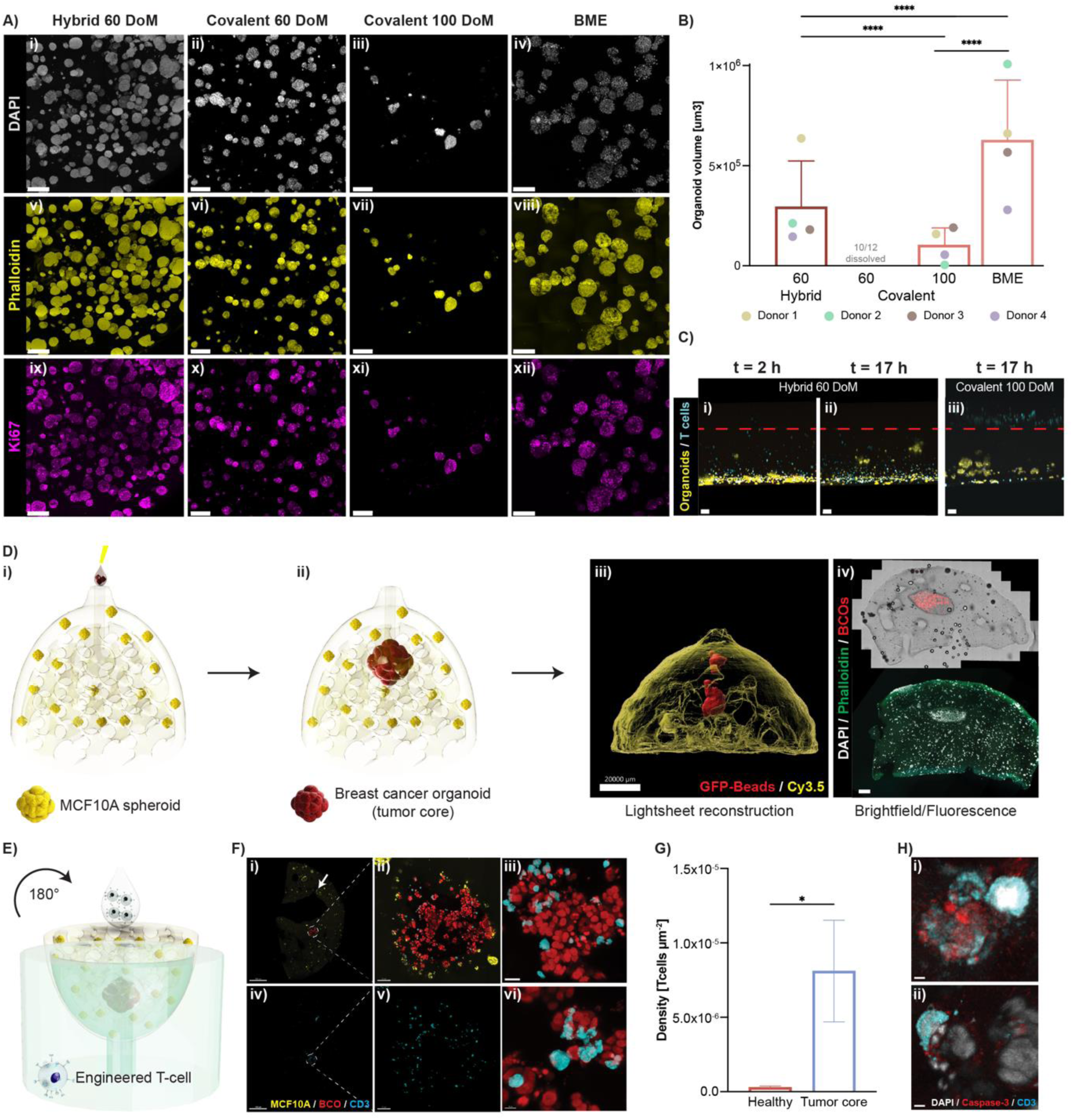
Hybrid gelMA is permissive for T cell targeting and migration. a) Representative immunofluorescent images of breast cancer organoids cultured in in hybrid 60 DoM (i,v,ix) and covalent 60 (ii,vi,x) and 100 (iii, vii, xi) DoM hydrogels and BME (iv, vii, xii) as a positive control. Scale bar = 200 µm. b) Corresponding quantification of organoid volume. N = 4. Each dot representing one donor. c) Side view of Tcell assay at t = 2 h and t = 17 h for hybrid 60 DoM gel (i, ii) stained for live (yellow) cells, Tcells (cyan) and for covalent 100 DoM gel at t = 17 h (iii). Scale bar = 50 µm. d) Schematic of volumetrically bioprinted gyroid breasts containing MCF10A spheroids and a pocket for tumor injection (i, ii). iii) 3D reconstruction of the printed breast (yellow) with injected fluorescent beads (red). Scale bar = 100 µm. iv) Confocal imaging showing the clearly from the rest of the breast distinguishable tumor core (red). e) Schematic of T cell exposure of bioprinted breasts. f) Confocal imaging showing T cell (cyan) infiltration into the breast containing MCF10A spheroids (yellow) and breast cancer organoids (red). i, iv) Whole breasts. Circle highlighting the tumor core. Arrow highlight a pore. ii, v) Zoom in into the tumor core. iii, vi) Zoom in into organoids that are interacting with T cells. g) Quantification of T cell densities in either the healthy or the tumor compartment obtained through 3D segmentation. h) BCO targeted by a T cell (cyan) and positive for cleaved Caspase-3.

### Hybrid hydrogels support cell-ECM interaction-driven mechanotransduction of hMSCs

Vascular network formation is crucially supported by MSCs through the secretion of pro-angiogenic factors, cell-cell contacts, matrix remodelling and mechanical stabilization of angiogenic sprouts^49–52^. In order to do so, sprouting endothelial cells recruit MSCs, which then migrate towards protrusions specifically protrusion tips or sprout roots to support their formation, elongation and maturation^53–55^. This pro-angiogenic capacity is regulated by stiffness-dependent mechanotransduction of the MSCs influencing their ability to spread, stretch and migrate through their microenvironment^56–58^.

We therefore wanted to see, if in accordance with better vascular network formation, MSCs would be able to migrate and spread through hybrid gel formulations. Accordingly, we cultured hMSCs in covalent 60 and 100 DoM formulations, as well as hybrid 60 DoM formulations, as this showed the best vascular network formation. Hybrid 60 DoM gel significantly outperformed both 60 DoM and 100 DoM covalent gels in terms of hMSC migration towards a chemoattractant (bFGF) over three days (**Fig. 3a,b**). This demonstrates the MSCs potential to break up reversible bonds. Further this could indicate that the enhanced network formation might be connected to improved ability of the cells to move and spread throughout their microenvironment. MSCs moreover displayed stretched morphologies in hybrid and covalent 60 DoM formulations as opposed to 100 DoM formulations. This was assessed through a decrease in circularity (**Fig. 3d,e**) and increase of cell area (**Extended Fig. 5a**). Furthermore, we observed an increase in nuclear area and decrease in nuclear circularity (**Extended Fig. 5b,c**). Similar results have previously been reported by Tang *et al.* for fast relaxing boronate-based hydrogels and were associated with improved cell-matrix interactions^59^.

We, therefore, sought out to have a closer look at cell-matrix interactions of hMSCs with our hybrid gel formulations. We stained MSC-gel samples for F-actin and paxillin, a focal adhesion protein involved in signaling between cells and the extracellular matrix. This revealed localization of paxillin along the membrane particularly focused on stretched cell regions, which also exhibited stress fibers, in both 60 DoM covalent and hybrid gels. In contrast paxillin stainings were diffuse, disorganized and homogeneous throughout the cell in 100 DoM hydrogels (**Fig. 3c**). This could be an additional indicator of improved cell-matrix interactions in hybrid and covalent 60 DoM materials. Since stretched cells and paxillin-rich adhesions are associated with increased activation of mechanotransduction pathways, we subsequently looked at the location of Yes-associated protein (YAP). This downstream actuator of paxillin has previously been associated with enhanced pro-angiogenic potential of hMSCs^56,60^. MSCs in covalent and hybrid 60 DoM gels displayed stretched morphologies and increased nuclear YAP translocation, particularly in those stretched cells, as compared to 100 DoM gels (**Fig. 3d-f, Extended Fig. 5a,d**). A significantly higher nuclear-to-cytoplasmic YAP ratio could be observed in hybrid gels compared to covalent formulations (**Fig. 3f**). However, despite the softer nature of 60 DoM gelMA, hybrid and covalent 60 DoM materials exhibited comparable proportions of stretched cells with nuclear YAP translocation (**Fig. 3g**). This can likely be attributed to the comparable stress relaxation kinetics of the two materials, which has been demonstrated fundamental in determining YAP translocation^61,62^. Mechanical stimulation and paxillin-driven translocation of YAP has not only been associated with increased pro-angiogenic capacity but also MSC fate determination, with nuclear YAP driving osteogenic gene expression^56,60,63–65^. We consequently investigated if osteogenic differentiation would be supported in our hydrogel formulations when additionally supplemented with osteogenesis promoting growth factors. As a proof-of-concept hMSCs were exposed to osteogenic differentiation medium for 21 days and Alizarin red staining to detect mineralization was performed. We detected comparable levels of mineralization in both covalent and hybrid 60 DoM materials, which was significantly higher than in covalent 100 DoM material (**Fig. 3h,i**). This indicates that both 60 DoM materials possess a similar capability of for supporting osteogenesis and that this capability could be partly steered by improved cell-matrix interactions and mechanotransduction-associated promotion of osteogenic gene expression.

### Supramolecular hybrid gels enable long-term organoid culture

Supramolecular gels have previously shown promising results for the culture of different types of organoids^21–23,25^. Importantly, matrigel, due to its unfavorable rheological properties, is not suited for many bioprinting approaches^26^. We, therefore, assessed whether we could culture organoids and combine organoid culture and volumetric bioprinting with our hybrid hydrogel formulations. We chose breast cancer organoids, as they are highly reliant on dynamic microenvironmental changes and due to the clinical relevance of this highly prevalent disease^66,67^.

Breast cancer organoids from four different donors were cultured in covalent 100 and 60 DoM and hybrid 60 DoM hydrogels, as well as the golden standard in organoid culture: BME. Notably, breast cancer organoids needed more time (∼ 7 days) to start proliferating after seeding when cultured in the covalent and hybrid gelMA hydrogels than in BME (∼ 3 days). This, is likely due to the fact that organoids require an extended recovery time after casting due to photoirradiation. Furthermore, organoids from all donors grew significantly smaller in different gelMA formulations than in BME, with the smallest and most sparse organoid growth occurring in covalent 100 DoM conditions (**Fig. 4a,b**). Three out of four organoid lines grew more densely (**Fig. 4A4a**) and formed proliferative organoids of a significantly increased volume in the hybrid gelMA than in the covalent 100 DoM gel (**Fig. 4A,B4a,b**). Notably, 60-100% of the gels made from covalent 60 DoM dissolved over the culture period. Images acquired for G60 were obtained from the few samples that did not dissolve in culture. As the bigger part of those samples was not stable in culture, it was excluded from further analysis. This is in line with our previous results, demonstrating impaired shape fidelity and long-term stability of covalent 60 DoM hydrogels compared to hybrid formulations.

Overall, organoids in all conditions presented with a high degree of Ki67 expression, indicating that they were in a proliferative state. Taken together, these results suggest, that while covalent 100 DoM hydrogels are stable over time in culture, they do not offer a permissive enough environment for organoid growth. Covalent 60 DoM hydrogels however allow for organoid growth, but are not stable over time. Hybrid 60 DoM hydrogel hereby exhibits a good balance of both permissiveness for organoid growth, as well as for long-time stability during prolonged culture (**Fig. 4A,B4a,b**).

### Hybrid gelMA is permissive to T cell migration

In a T cell co-culture assay previously developed by our group, T cell behavior patterns when exposed to breast cancer organoids could be identified via live imaging^68^. However, in this setting T cells were added to a breast cancer organoid suspension containing low concentrations of BME. This facilitates T cell movement, not taking into account tumor infiltration as one of the main challenges for cellular immunotherapies in the treatment of solid tumors. Here, this assay was adapted to be executed with hydrogel formulations to examine, whether T cells would be able to sense breast cancer organoids and migrate into the solid gel. Engineered T cells (TEGs^69–71^) were added on top of the preformed BCO-laden gels (250 – 300 µm thickness) and live imaging was performed for ∼17h (**Extended Fig.6**). T cells were able to migrate through the hybrid gel to reach the organoids already within the 2 hours of preparation time confocal imaging (**Fig. 4c, i, ii**). In covalent 100 DoM however, T cells had not yet infiltrated the gel after 17 hours (**Fig. 4c, iii**).

Next, we wanted to test whether T cell infiltration can be targeted to tumor organoids within larger and more physiologically-relevant structures and scales. Hence, we volumetrically bioprinted a breast shaped model with healthy spheroids (MCF10A) embedded in the gel. BCOs were injected into a 3.75 mm diameter pocket within the breast (**Fig. 4di, ii**). BCOs grew within the pocket and remained constricted to the tumor pocket for the duration of culture (1 week) and were clearly distinguishable from normal spheroids (**Fig. 4Diii4diii, iv**). We added T cells to the bottom side of the mini breasts, which we flipped upside down and inserted into a custom PDMS holder. Like this, they could enter the pores of the gyroid-shaped model (**Fig. 4e**). Due to the large distance (3.9 mm from the bottom of the breast until the nearest point of the tumor pocket) T cell infiltration took longer to reach tumor organoids than in the previous flat and thin samples. However, T cells successfully infiltrated the tumor regions through the pores (**Fig. 4F4f**) within three days. Cross-sections of the mini breasts show that T cells were almost exclusively located within the tumor pocket (**Fig. 4f, g**) and directly interacting with the BCOs (**Fig. 4fiii, vi**) or the pores. BCOs further stained positively for cleaved Caspase-3 (**Fig. 4h**). These data indicate that T cells not only can migrate long distances within hybrid 60 DoM hydrogels but can also specifically target BCOs. Bioprinted breast models from hybrid hydrogel could thus be a promising in vitro platform to study targeted immunotherapy for solid tumors.

## Conclusions

In this study, we introduced formulations composed of adamantane isothiocyanate-modified gelMA and acrylated beta cyclodextrin to produce supramolecular hybrid bioresins. We demonstrated, that cyclodextrin and adamantane form dynamic and reversible host-guest interactions. Those cause an increased material stiffness, denser polymer network formation, slower degradation kinetics and enable viscoelastic behavior of our hybrid gel formulations. We showcased the suitability of our hybrid formulations for volumetric bioprinting. Hybrid hydrogels prints yielded excellent shape fidelity even at low DoM that are not capable of forming self-suporting structures in covalent gel formulations with the same DoM. Additionally to improved long-term stability and shape fidelity, we demonstrated the permissiveness of the supramolecular bonds for cellular growth, migration and remodeling. This was shown for different cell types including endothelial cells, MSCs and breast cancer organoids. Hybrid formulations were superior to covalent formulations for vascular network formation and enabled MSC migration and mechanotransduction. Further, they displayed similar potential for osteogenic differentiation as covalent gels with the same DoM. Furthermore, we demonstrated the compatibility of hybrid formulations with organoid culturing and extensive T cell migration in large-scale bioprinted construct. Taken together our results demonstrate that supramolecular hybrid gelMA provides an excellent balance between mechanical demands (shape fidelity) and permissiveness for dynamic cellular processes, which is a crucial challenge in the development of biomaterials for tissue engineering.

## Acknowledgements

This project received funding from the European Research Council (ERC) under the European Union’s Horizon 2020 research and innovation program (grant agreement no. 949 806, VOLUME-BIO). R.L and J.M. acknowledge the funding from the Gravitation program “Materials Driven Regeneration” (023.003.013) funded by the Netherlands Organization for Scientific Research (024.003.013). R.L. acknowledges financial support from Dutch Research Council’s Talent program (Vidi, 20387).

## Disclosures

M.F, P.N.B, M.B, A.R., and R.L. are inventors on a provisional patent application that covers the hydrogel reported in this manuscript and its application for bioprinting, and cell and organoid culture. R.L. is scientific advisor for Readily3D SA. The other authors declare no competing interests.

---

**Extended Figure 1.**
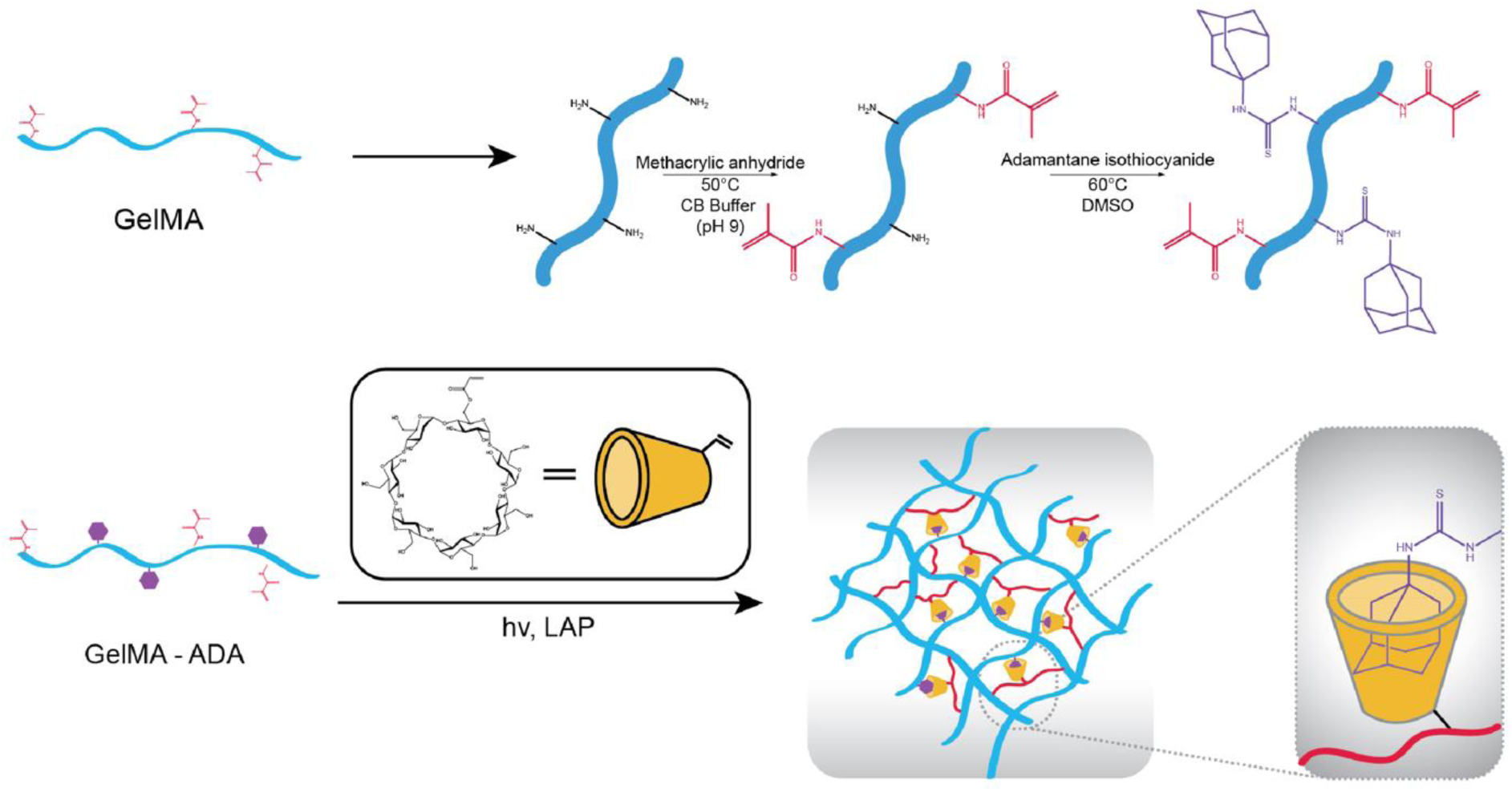
Schematic of gelatin functionalization with methacrylic anhydride and adamantane isothiocyanide as well as schematic illustration of host guest interactions formed between adamantane groups and cyclodextrin.

**Extended Figure 2.**
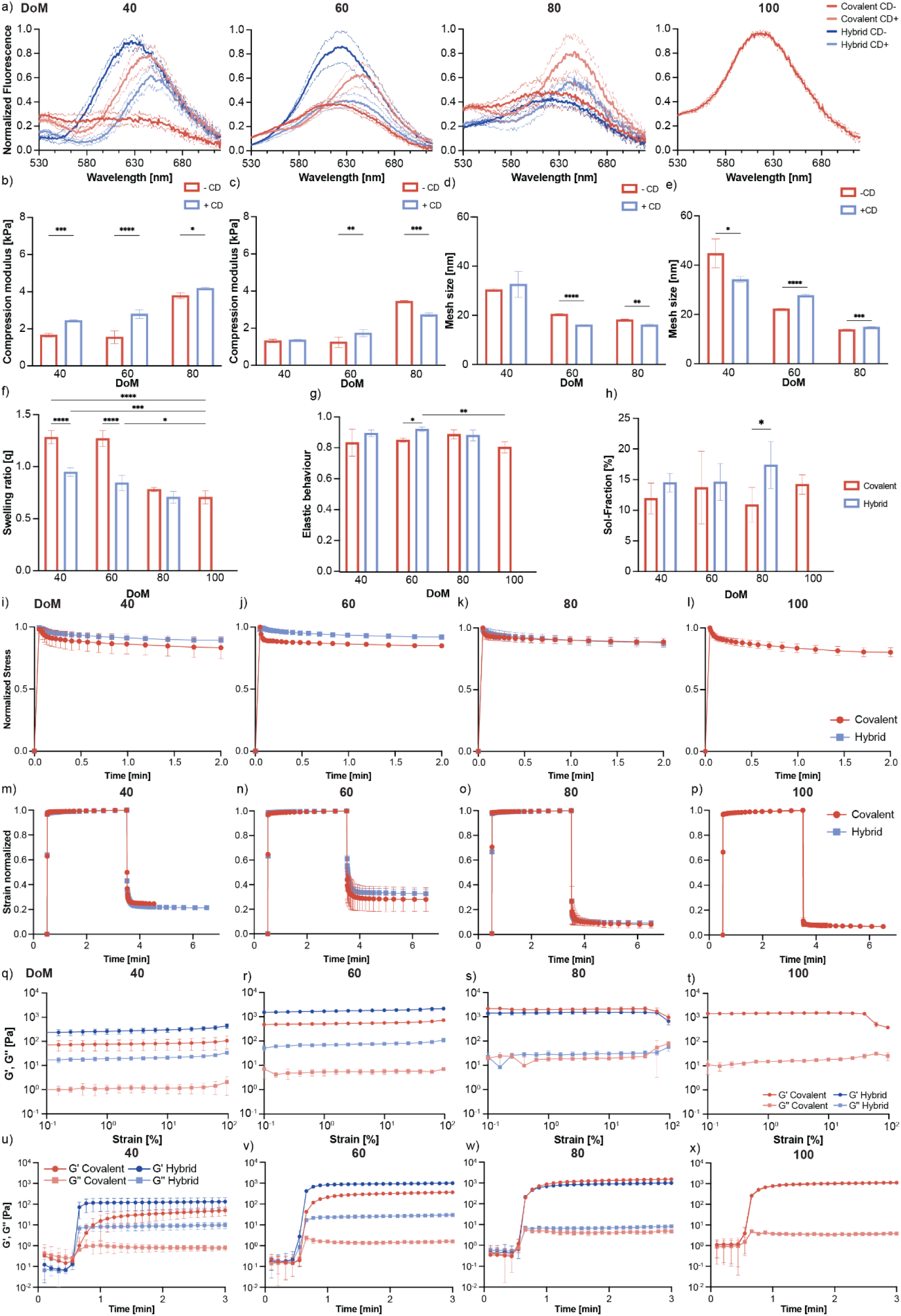
Mechanical and physical characterization of 5 w/v% covalent and hybrid bioresins with varying DoM (40, 60, 80, and 100). a) Nile red assay results indicating hydrophobicity as a function of fluorescence with and without addition of cyclodextrin (CD). Compression modulus of hybrid (b) and covalent (c) formulations with and without cyclodextrin. Mesh sizes of hybrid (d) and covalent (e) formulations with and without cyclodextrin. f) Swelling ratio, g) Elastic behavior and h) sol fraction of covalent and hybrid formulations of different DoM. Stress relaxation kinettics (i-l), creep (m-p), amplitude sweep (q-t) and photorheology (u-x) of covalent and hybrid formulations of different DoM.

**Extended Figure 3.**
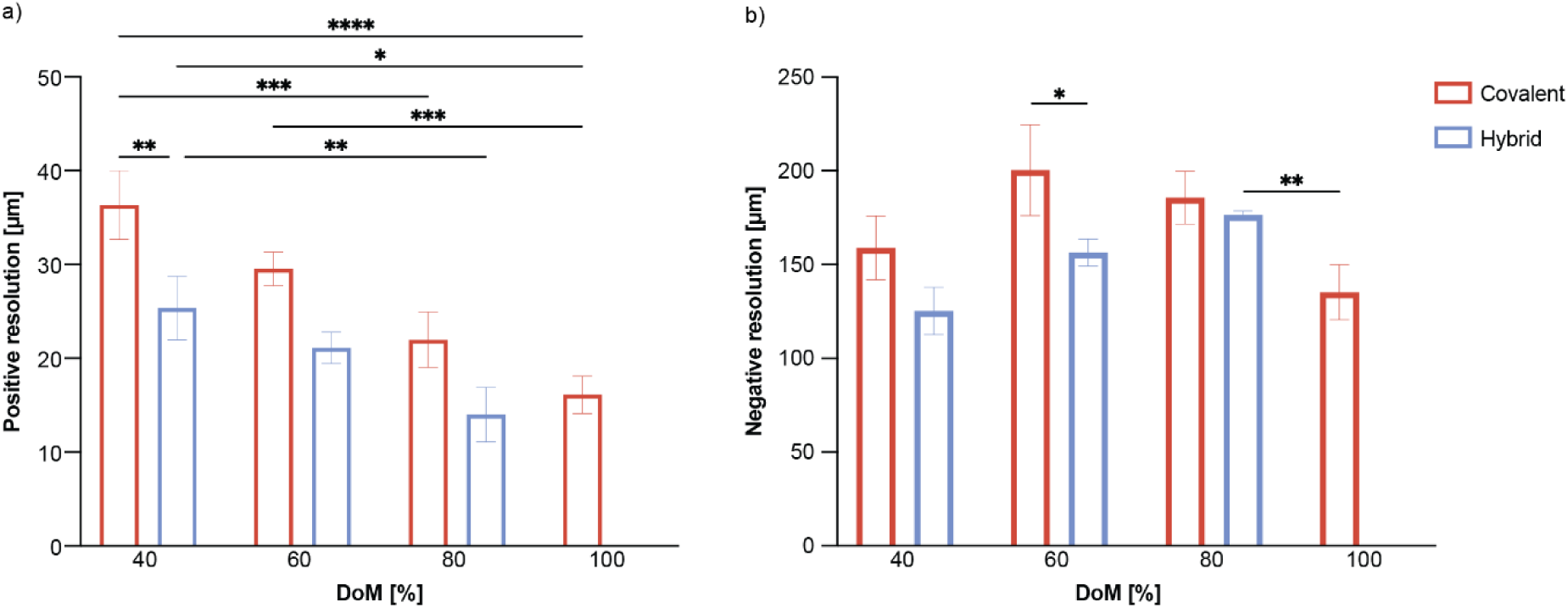
Volumetric printing resolution of positive (a) and negative (b) features for covalent and hybrid formulations.

**Extended Figure 4.**
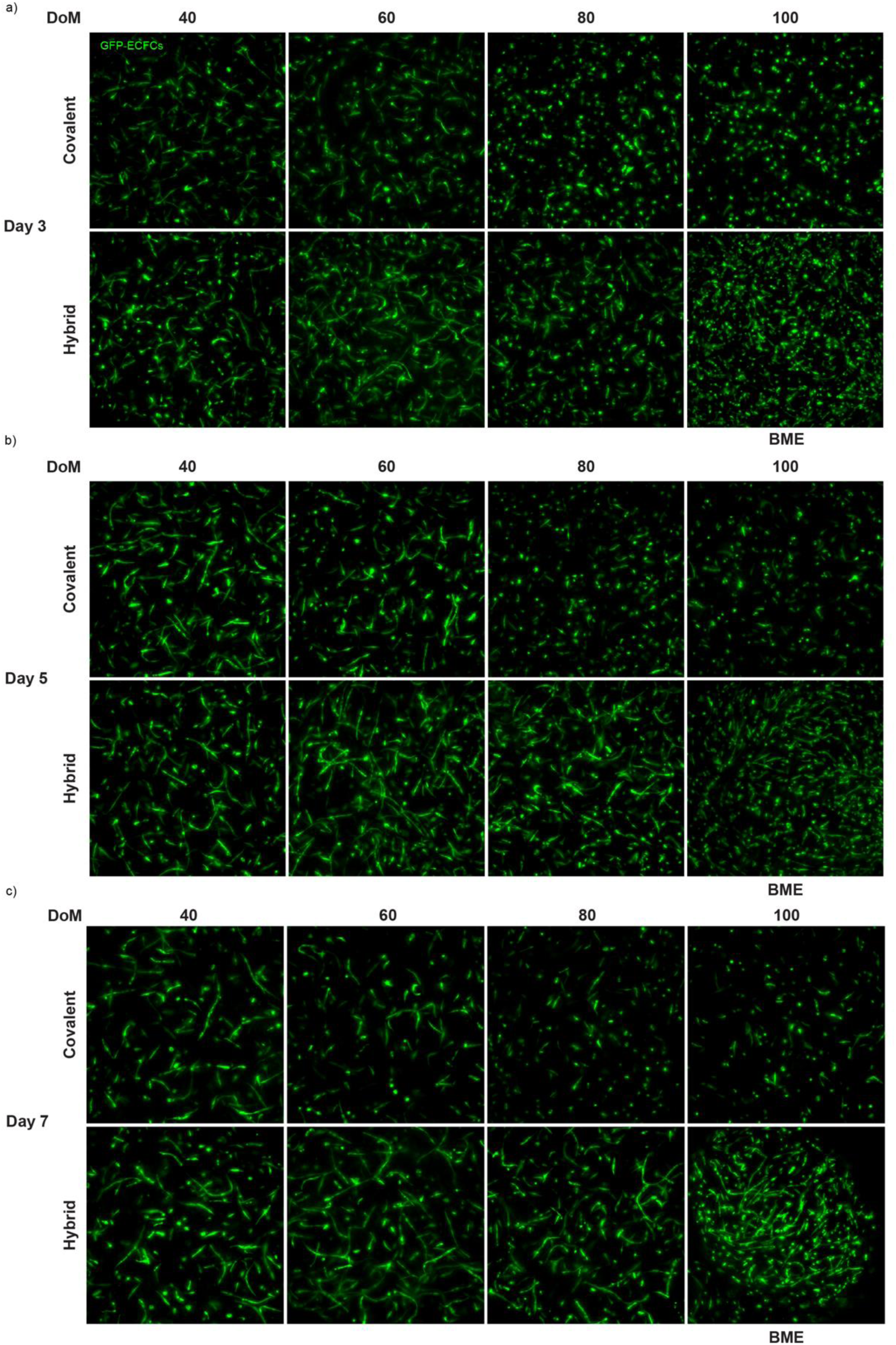
Vascular network formation of ECFCs in covalent and hybrid formulations of different DoM at day three (a), five (b) and seven (c).

**Extended Figure 5.**
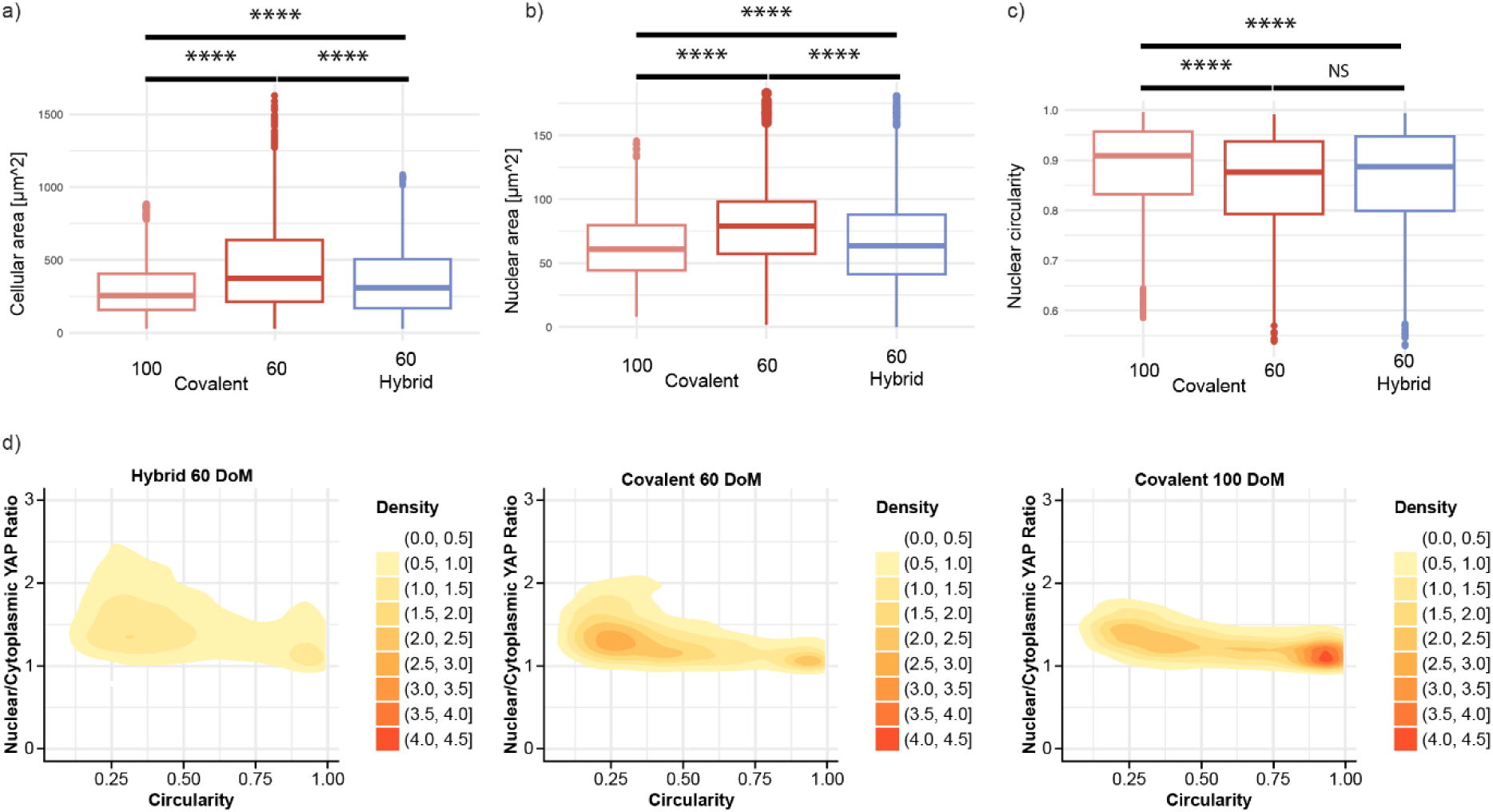
Characterization of cell morphology. Cellular area (a), nuclear area (b) and nuclear circularity (c) of MSCs cultured in 60 DoM hybrid and covalent formulations, as well as in 100 DoM formulations for three days. d) Density distribution of cells of different nuclear/cytoplasmic YAP ratios against cell circularity in samples cultured in hybrid 60 DoM and covalent 60 and 100 DoM hydrogels after 3 days.

**Extended Figure 6.**
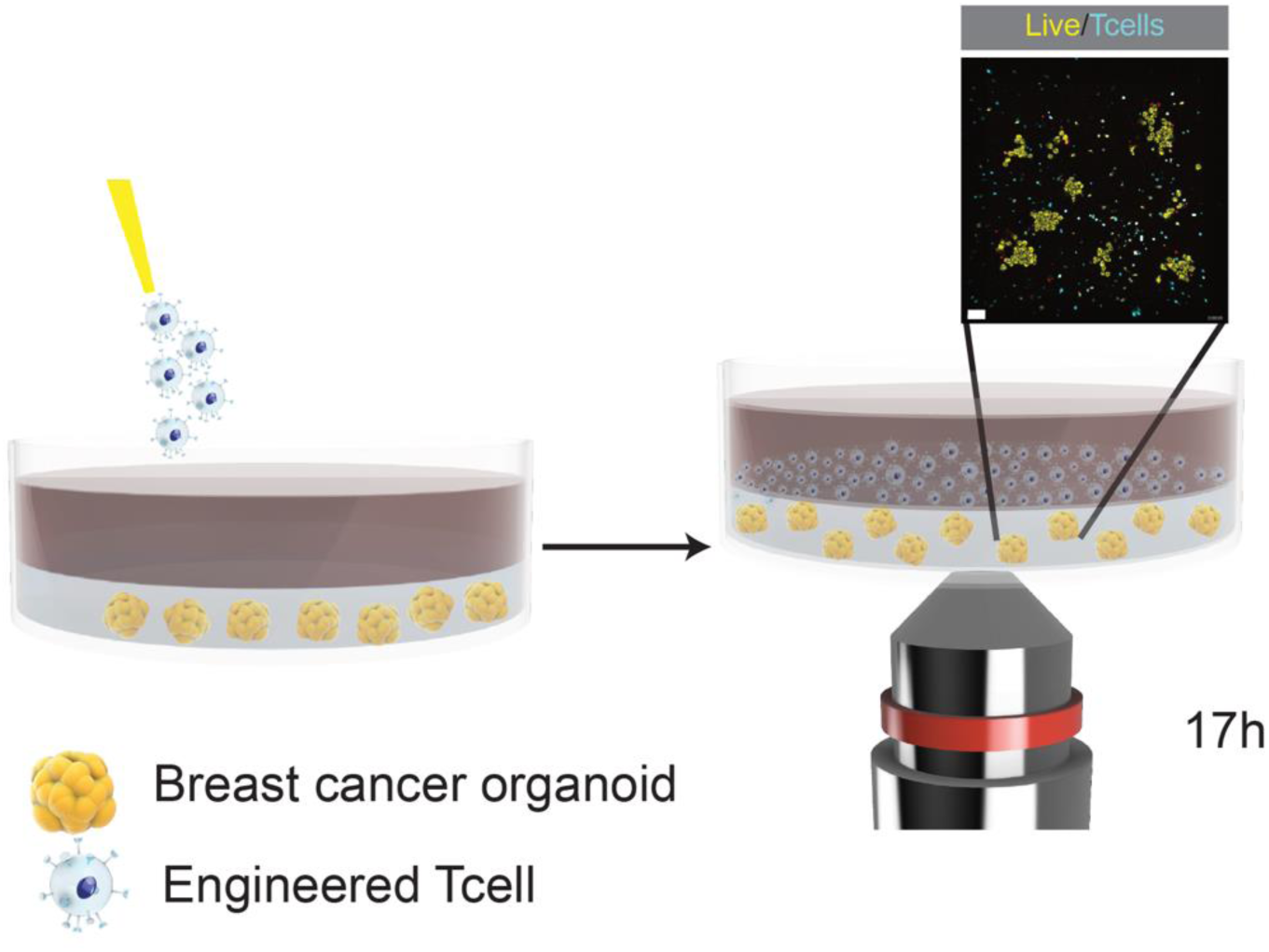
Schematic illustration of T cell pilot assays.

## Materials and methods

### Materials

Cy3-PEG-SH (Mw = 5 kDa) was purchased from Biopharma PEG (Watertown, USA). PDMS was purchased from Dow (Dow, SYLGARD™ 182 Elastomer kit). All other chemicals or reagents were purchased from Sigma-Aldrich unless stated otherwise.

#### Covalent gelMA synthesis

Type A porcine gelatin (Sigma) was dissolved in a CB-Buffer (carbonate-bicarbonate buffer pH 9, 0.1 M concentration) at 50°C at a final concentration of 10 w/v%. A constant temperature of 50°C was maintained throughout the synthesis process. 0.71 µL of methacrylate anhydride (MAA) per gram of gelatin per degree of methacrylation (DoM) was used to reach different DoMs. MAA was added 6 times at 11 minute intervals. The pH was stahilized at a pH of 9.0 using 5 M NaOH after each addition of MAA. The reaction was stopped after 90 minutes by lowering the pH to 7.4 using 1M HCl. The solution was centrifuged at room temperature for 5 min at 4000 rpm. The supernatant was collected and diluted to a final concentration of 5% w/v and then dialyzed against MilliQ water for 4 days at 4°C using cellulose dialysis membrane tubes (molecular weight cutoff = 12 kDa; Sigma-Aldrich). After dialysis, the gel was liquified by heating it to 50 °C and subsequently diluted with MilliQ to a final concentration of 2.5% w/v and sterile filtered. Following the sterile filtration, the solution was frozen to −80°C and lyophilized.

### Hybrid gelMA synthesis

GelMA was dissolved in DMSO at a final concentration of 10 % w/v and heated to 60 °C. The temperature was kept stable throughout the synthesis process. 5 mg of adamantane isothiocyanate (AITC; Sigma-Aldrich) per gram of gelMA per degree of supramolecular modification (DoSM) were added to the solution and stirred for 5 hours. The solution was cooled to room temperature to stop the reaction. Subsequently, the solution was centrifuged at room temperature for 5 minutes at 4000 rpm. The supernatant was dialyzed against MilliQ water for 4 days at 4°C resulting in the solution turning white. Afterwards, the solution was heated to 50°C, diluted with MilliQ to a final concentration of 2.5% w/v and centrifuged at room temperature for 5 minutes at 4000 rpm. The supernatant was heated to 50°C and sterile filtered. Following sterile filtration, the solution was frozen to −80°C and lyophilized.

### Acrylated-β-cyclodextrin (Ac-β-CD) synthesis

**Extended Figure 7.**
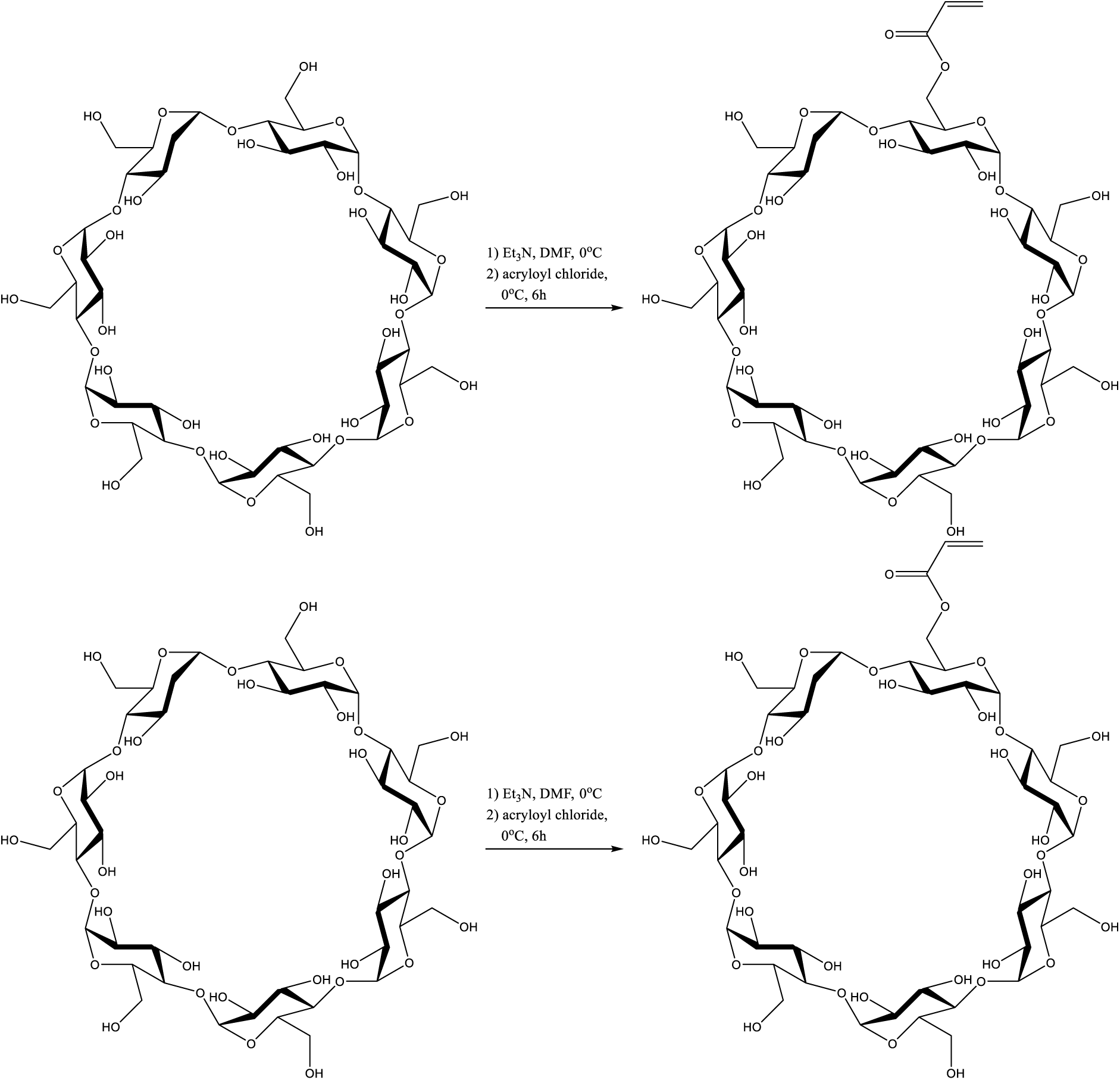
Ac-β-CD synthesis.

Ac-β-CD was synthesized as previously described^72^. Briefly, beta Cyclodextrin (β-CD; Sigma-Aldirch) (1.0 eq) was added to a mixture of Et_3_N (5.7 eq) in DMF (220 eq). The mixture cooled to 0°C under stirring, after which acryloyl chloride (7 eq) was added. The reaction was stirred for 6 h at 0°C. The mixture was filtrated to remove trimethylamine hydrochloride. Subsequently, the solution was concentrated using rotary evaporation and added dropwise to acetone at 0°C (900 eq) to allow the modified cyclodextrin to precipitate. The precipitate was washed with acetone and dried in vacuum for one day.

### Hydrogel precursor preparation and photocrosslinking

Stock solutions were prepared as follows: Covalent or hybrid gelMA were dissolved in PBS at a concentration of 10% w/v. Lithium phenyl-2,4,6-trimethylbenzoylphosphinate (LAP, Tokyo Chemical Industry) was dissolved in PBS at a final concentration of 1 w/v%. Acrylated-β-cyclodextrin dissolved in PBS at a final concentration of 35 mM. All stock solutions were heated to 37 °C until completely dissolved. Stock solutions were combined to make hydrogel precursor formulations at a final concentration of 5% w/v gelatin-based material, 0.1% w/v LAP, and an equimolar amount of adamantane to cyclodextrin, which was dependent on the DoSM of the corresponding material. All hydrogel formulations were prepared at these concentrations, with constant concentration of LAP and ratio of adamantane to cyclodextrin unless stated otherwise. The hydrogel formulations were crosslinked using a UV-oven (Cl-1000, Ultraviolet Crosslinker, λ = 365 nm, I = 8 mW/cm2 UVP, USA) unless stated otherwise.

### Volumetric printing

Hydrogel solutions were dispensed into cylindrical borosilicate glass vials (Ø 10 mm). Vials were loaded into a commercial volumetric 3D printer (Tomolite V2, Readily3D, Switzerland; 405 nm; 11.98 mW cm-2 light intensity). The samples were thermally gelated at 4°C prior to printing. STL files were loaded into the printer software (Apparite, Readily3D, Switzerland). After printing, the vials were heated to 37°C and printed constructs were washed gently with 37 °C PBS to retrieve the prints. Samples were submerged in 0.1% w/v LAP solution and irradiated for 1 min in a UV oven to ensure homogenous cross-linking,

### Measurement of the compression modulus via DMA

Cylindrical hydrogel samples (Ø 6 mm, 2 mm height) were casted by crosslinking for 10 minutes within a mold. To reach equilibrium swelling, samples were washed in PBS at 37°C overnight. Using a dynamic mechanical analyzer (DMA Q800, TA Instruments, The Netherlands) a strain ramp at 20% min-1 strain rate was applied until 30% deformation to assess the compressive properties, The compression modulus was calculated as the slope of the stress/strain curve in the 10–15% linear strain range.

### Measurement of stress relaxation and creep

Samples were subjected to a strain recovery measurement at a constant 20% strain for 2 minutes and left for recovery for 1 minute, with a preload force of 0.0010 N to assess the viscoelastic properties. The elasticity index was determined as the ratio between the recovered stress and the maximal stress under constant strain. Creep measurements were taken at a constant stress of 2 kPa for 3 minutes and left for recovery for 3 minutes. The obtained strain measurements were normalized for every sample.

### Photorheology, Frequency sweep, amplitude sweep, Tan δ

Using a DHR2 rheometer (TA Instruments, The Netherlands), crosslinking kinetics were assessed on hydrogel precursor solutions. Time sweep experiments were performed at a frequency of 1.0 Hz, angular frequency of 6.283 rad/s, with 1.0% constant strain at 21°C (n = 3 independent samples). The light source was activated (1200mha, AOMEES, China, λ = 365 nm, intensity of 24 mW/cm2 for the remaining 2.5 minutes) 30 seconds after the start of the measurement. Freguency sweep experiments were performed at a frequency range from 0.1 Hz to 100 Hz with 1.0% constant strain at 21°C. Subsequently, amplitude sweep experiments were performed at a frequency of 1.0 Hz at a strain rate from 0.1% to 100% strain at 21°C. 100 µL of gel were used with a gap size of 300 µm. A 20.0 mm parallel EHP stainless steel plate was used as geometry. Tan δ was calculated as the ratio of the storage and loss modulus taken from the frequency sweep measurements.

### Mesh size determination

Hydrogel mesh size was calculated through G’^max^ from frequency sweep measurements using **Formula 1** as an estimation of the mesh size as previously described^73^.

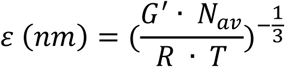

**Formula 1 Mesh size formula for analysis of hydrogels.**

### Macromer concentration, Sol-fraction, and Swelling ratio

The macromer concentration, sol-fraction, and swelling ratio experiment was performed according to a recent publication^37^. Briefly, crosslinked cylindrical hydrogels samples were weighed immediately after crosslinking to determine their initial mass. After incubation in PBS at 37°C over night the samples were weighed again, and their mass was measured as mass_wet,t0_. Subsequently the hydrogels were lyophilized, and the dry mass (mass_dry,t0_) was measured. The samples were stored in PBS again to ensure swelling of the dry gels and placed in the incubator at 37°C overnight. The wet mass of the hydrogels was measured as mass_wet,t1_. The samples were lyophilized, and the mass of the dry samples was measured as mass_dry,t1_. The macromer concentration of the hydrogel formulations was calculated with the following formula:

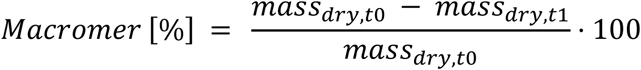

**Formula 2 Formula to calculate the macromer concentration.**

The sol-fraction of the hydrogel formulations was calculated with the following formula:

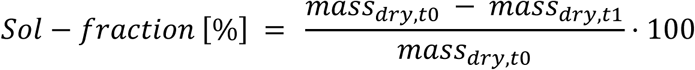

**Formula 3 Formula to analyse the crosslinking properties of different gelMA formulations via the Sol-fraction.**

The swelling ratio of the hydrogel formulations was calculated with the following formula:

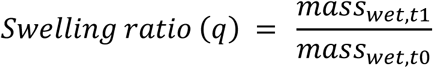

**Formula 4 Formula to calculate the swelling ratio.**

### Nile-Red

Wells of a 96-multiwell plate were filled with 12 µL of a solution of Nile-Red dye in methanol at a concentration of 0.005 mg/mL. The methanol was allowed to evaporate overnight. 200 µL of hydrogel precursor at a concentration of 0.5 % w/v gelMA and 0.01 w/v% LAP with an equimolar ratio of adamantane to cyclodextrin, were added, UV exposed for 1 minute and equilibrated overnight. 200 µL MilliQ were measured as a negative control. The obtained data was normalized for every sample.

Enzymatic degradation assay.

Hydrogel samples were allowed to swell over night in PBS. Subsequently they were incubated in 0.2% w/v collagenase type II in Dulbecco’s modified Eagle Medium (DMEM, 31966, Gibco, The Netherlands) supplemented with 10% v/v heat-inactivated fetal bovine serum (FBS Gibco, The Netherlands), and 1 % v/v penicillin and streptomycin (Life Technologies, The Netherlands) at 37°C. At different time points (15, 30, 45, and 60 minutes, n = 3 independent samples per time point) samples were removed from the enzymatic solution and the mass was measured and compared to the initial hydrogel mass to determine the degradation rate of samples over time.

### Shape fidelity

Hollow cylindrical models (Ø 6 mm, 2 mm height; hole Ø 3 mm) were printed from different hydrogel formulations using the volumetric bioprinter, washed in warm PBS and postcured in a 0.1% w/v LAP solution for 1 minute in the UV-oven. To assess shape fidelity, hollow cylinders were oriented onto a glass slide on their sides and images of the open hollow structures were taken. Shape fidelity was calculated by dividing the height over the width of the inner hollow cylinder. These calculations were then normalized per sample.

### Cell culture

#### MSCs

Human MSCs were isolated from bone marrow aspirates as previously described^74^ from patients giving informed consent and as approved by the research ethics committee of the University Medical Center Utrecht. MSCs were expanded in αMEM (22 561, Invitrogen, Carlsbad, USA) supplemented with 10% heat-inactivated fetal bovine serum, 0.2 mm L-ascorbic acid 2-phosphate (A8960, Sigma-Aldrich, St. Louis, USA), 100 U mL^−1^ penicillin with 100 mg mL^−1^ streptomycin (15 140, Invitrogen), and 1 ng mL^−1^ basic fibroblast growth factor (233-FB; R&D Systems, Minneapolis, USA).

#### ECFCs

ECFCs were isolated from human cord blood from consenting mothers as previously described^74^ and as approved by the medical research ethics committee of the University Medical Center Utrecht. ECFCs were expanded in EGM-2 Endothelial Cell Growth Medium-2 BulletKit (Lonza, The Netherlands) supplemented with 10% FBS.

#### MCF10A

MCF10A normal epithelial cells (ATCC CRL-10317 ™) were cultured as to manufacturers recommendations. Briefly, MCF10A were seeded in T75 culture flasks and MCF10A medium (Advanced DMEM + 5% v/v Horse serum + 1% v/v P/S + 20 ng/µL EGF + 0.5 µg/mL Hydrocortisone + 10 µM Forskolin + 10 µg/mL Human insulin (Sigma 11061-68-0)) was added and refreshed once a week. Once confluent cells were split at a 1:6 ratio.

#### Breast cancer organoids

Breast cancer organoids (BCOs) were cultured as described previously^75^. Briefly, organoids were resuspended in the appropriate volume of cold basement membrane extract (Basement Membrane Extract, Cultrex RGF BME type 2, R&D systems, 3533-005-02) and plated as small drops in suspension culture plates (100 µL per well of a 12 multiwell plate). The BME was allowed to solidify at 37°C for 10 minutes. 750 µL of type 1 expansion medium^76^ (Extended Table 1) from now on referred to as BCM were added per well. Medium was refreshed twice a week. Breast cancer lines were passaged once a week at indicated splitting ratios and TryplE (TrypLE Express Enzyme, Trypsin replacement with EDTA, Thermo Fisher, 12605-010) concentrations (Extended Table 2).

**Extended Table 1.**
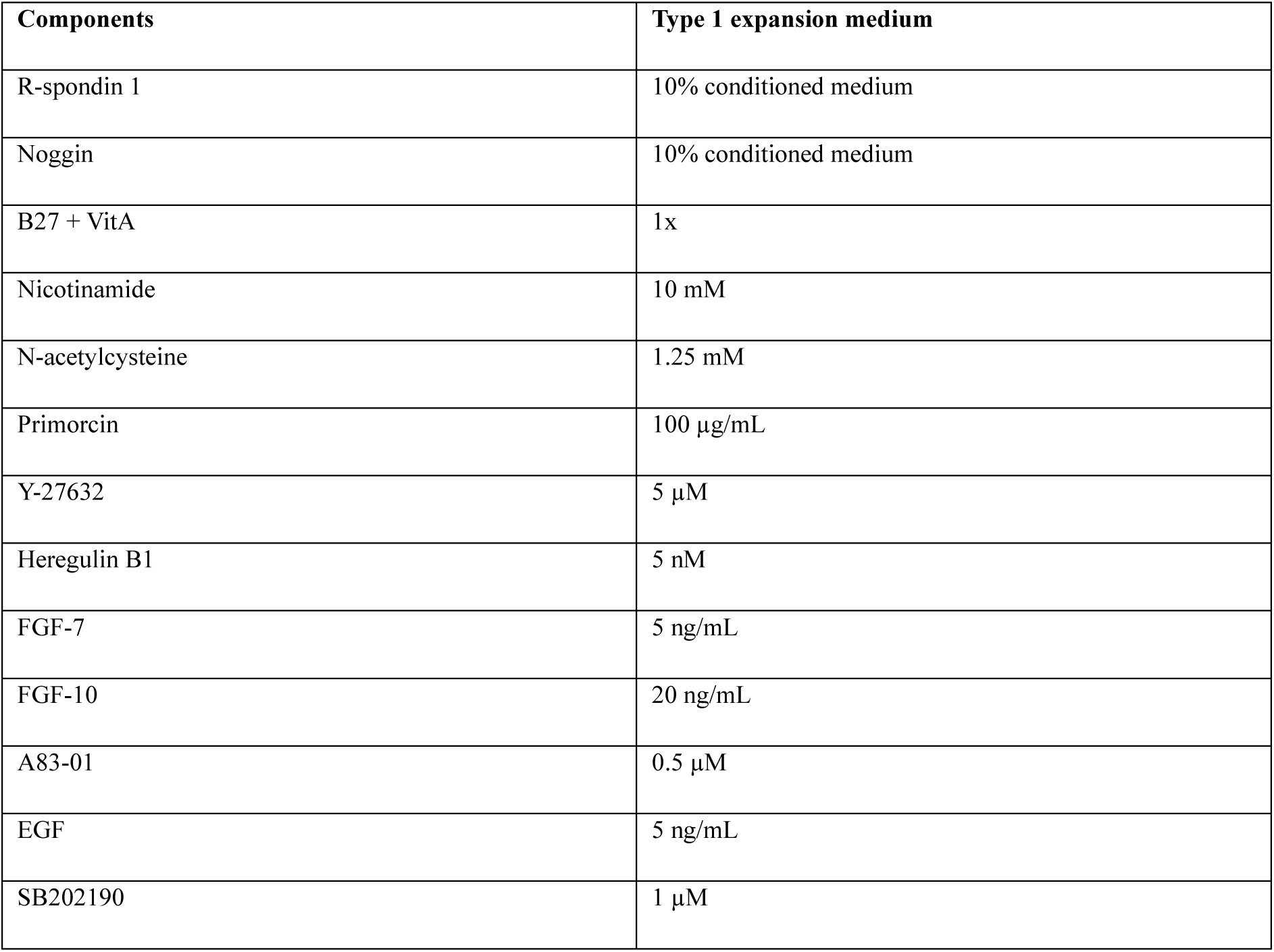
Type 1 medium composition in F12+++ (Advanced DMEM F12 (Gibco, 12634-010) supplemented with 1 % v/v HEPES, 1 % v/v Glutamax and 1 % v/v penicillin–streptomycin (P/S))

| Components | Type 1 expansion medium |
| --- | --- |
| R-spondin 1 | 10% conditioned medium |
| Noggin | 10% conditioned medium |
| B27 + VitA | 1x |
| Nicotinamide | 10 mM |
| N-acetylcysteine | 1.25 mM |
| Primorcin | 100 μg/mL |
| Y-27632 | 5 μM |
| Heregulin B1 | 5 nM |
| FGF-7 | 5 ng/mL |
| FGF-10 | 20 ng/mL |
| A83-01 | 0.5 μM |
| EGF | 5 ng/mL |
| SB202190 | 1 μM |

**Extended Table 2.**
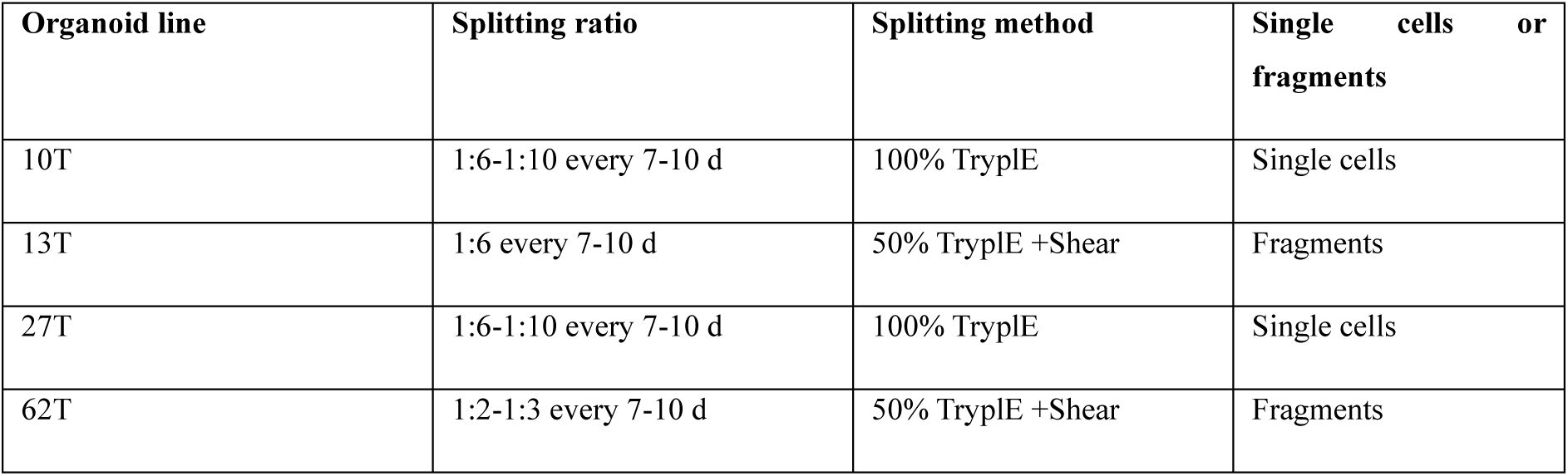
Organoid culturing guidelines.

### Vascular network formation assay

The outer ring of the clover molds was filled with 10% w/v 100 DoM gelMA and crosslinked under UV exposure for 1 minute. ECFCs and MSCs were embedded in covalent and hybrid gels, as well as BME at a ratio of 1:3 and a final density of 7.5 million cells per mL, casted into the cylindrical inlets of the clover mold, and UV exposed for 8 minutes. Samples were cultured in EGM-2 media + 10 % v/v FBS for one week. Media was refreshed every other day and samples were imaged with a THUNDER imaging system (Leica) on day 1, 3, 5 and 7. The software AngioTool^77^ was used to evaluate vascular network features.

### MSC migration assay

The middle channel of chemotaxis µ-slides (Ibidi) was filled with 10 µL of either 60 or 100 DoM covalent gel or hybrid 60 DoM gel and crosslinked under UV exposure for 2 minutes. 3 million DiD labelled hMSCs per mL in MSC media were injected into one of the media chambers, according to manufacturer’s instruction. The other chamber was filled with MSC media containing 5ng/mL bFGF. Chambers were cultured for 3 days and media was refreshed daily. Migrating cells were imaged with a THUNDER imaging system (Leica) after fluorescent labeling with Did tracking dye (Vybrant DiD Cell-Labling Solution, Thermo Fisher Scientific).

### Osteogenic Differentiation

MSCs were differentiated in a commercial osteogenic differentiation medium (Human Mesenchymal Stem Cell (hMSC) Osteogenic Differentiation Medium BulletKit) for 21 days. Samples were fixed in 4% w/v PFA for 1 h at room temperature and, after dehydration in a series of increasing ethanol solutions (70–100%), were cleared in xylene. Subsequently, the samples were embedded in paraffin and sliced into 5 µm thick sections (Microm HM340E; Thermo Fischer Scientific). Prior to staining, tissue sections were deparaffinized with xylene and gradually rehydrated through decreasing ethanol solutions (100–70%). Alizarin red solution was added for 2 minutes. Sections were dehydrated in acetone, acetone-xylene and xylene. Slides were imaged were with a THUNDER imaging system (Leica). Mineralized areas (red) were expressed as a percentage of the total hydrogel volume.

### Adipogenic differentiation

Adipogenic differentiation was induced by culturing MSCs in adipogenic differentiation medium made from αMEM containing 10 % v/v FCS, 100 U/mL penicillin, 100 μg/mL streptomycin, 0.4 μg mL−1 dexamethasone (Sigma–Aldrich, The Netherlands), 0.052 mm 3-isobutyl-1-methylxanthine (IBMX; Sigma–Aldrich, The Netherlands), 0.2 mm indomethacin and 10 µg/mL insulin (Sigma–Aldrich, The Netherlands) and medium was refreshed every twice a week.

Samples were fixed in ice cold 4% w/v PFA for 1 h at 4 °C and permeabilized in 0.1% Triton-100 for 15 minutes at room tempereature. Samples were blocked by washing three times for 5 min with a 3% Bovine Serum Albumin (BSA, Sigma) solution in PBS. Subsequently, gels were stained for lipid vesicles using LipidTox (1:200; Invitrogen, HCS LipidTOX Green Neutral Lipid Stain) and DAPI (1:1000) for 3 h at room temperature in the dark. Samples were washed with PBS and kept in the dark until image acquisition.

### Hydrogel discs for breast cancer organoid culture

Using FDM printing of PLA a mold was printed (Suppl. Fig.8A). The mold was filled with PDMS (Dow, SYLGARD™ 182 Elastomer kit) to make a two part (Top & Bottom) mold for hydrogel discs (9 discs a 3 x1.5 mm per disc). Top parts were glued to bottom parts using a thin layer of 10% w/v 100 DoMg. 6% w/v gelMA formulations with 0.1% w/v LAP were prepared. For supramolecular gelMA equimolat amounts of Ac-β-CD were added.

Organoids were fragmented in TryplE for 5-15 minutes. Fragments were collected in cold F+++ and spun down at 300 g for 5 min at 4°C. Pellets were washed with PBS + 0.1% BSA divided over four Eppendorf tubes and spun down again. Pellets were resuspended in 150 µL hydrogel and 15 µL gel were added per well of the mold. Molds were covered with glass slides to exclude oxygen from the reaction. The filled molds were exposed to UV light for 2 minutes in the UV oven. Discs were removed from the mold using a spatula, added into one 48 multiwell plate well per disc and covered with 100 µL of BCM. As a control, BCOs were plated in drops of 15 µL BME. Medium was refreshed twice a week. Discs and controls were cultured for 10-14 days.

Stainings were performed as previously described^78^. Briefly, discs were fixed in ice cold 4% w/v PFA for 45 min on a rocker at 4°C. Samples were washed with PBT (PBS + 0.1% w/v Tween20) and kept in PBT until further processed. Before staining, samples were washed in wash buffer 2 (WB2; 1 L PBS + 1 mL of Triton X-100 + 2 mL of 10% (w/v) SDS + 2 g BSA) for 30 min at 4 °C. Samples were stained for Ki67 (eBioscience 14-5698-82; 1:200) in WB2 overnight. The next day, samples were washed 3x for 1.5 h in WB2. Samples were stained for DAPI (1:1000; Thermo Fisher Scientific Invitrogen™ D1306), Phalloidin-Atto700 (1:200; SigmaAldrich 79286-10NMOL) and donkey-anti-mouse-AF647 (1:200, ThermoFisher Scientific A31570) over night. The next day samples were washed 3x 1.5 h in WB2 and mounted onto thin glass slides. A z-stack of 150-200 µm size was aquired at 20X magnification per sample (triplicates). Organoids were segmented using Imaris (Oxford Instruments) based on the phallolidin signal and the volume of single organoids was extracted.

### T cell pilot experiments

60 DoM hybdrid hydrogels (6 \% w/v) were prepared as described above. BCO donor 13T was collected in ice cold F+++ and filtered over a 70 µm strainer. Organoids were spun down at 300 g for 5 min at 4 °C. Pellets were resuspended in PBS+0.1 % BSA stained with Cell tracker green (Cell tracker green Invitrogen™ C12881; 1:1000) for 15 min, washed with PBS + 0.1 % w/v BSA and spun down again. Pellets were resuspended in 100 µL of hydrogel and 15 µL per well were plated. Under hypoxia, gels were crosslinked using a UV lamp for 2 minutes. As a control, some organoids were plated in 100% BME and some in suspension containing 2.5% BME as previously described^79^. Organoids were allowed to rest for three days and media was refreshed on day three. On day four after seeding, engineered T cells (TEGs) were labelled with eFluor 450 (eBioscience Cell Proliferation Dye eFluor 450 Thermo Fisher; 1:3000) for 10 min, washed with PBS and then added on top of the gels (10 k per well). 40 µL imaging medium (1:1 BCM:T cell medium (RPMI-GlutaMAX supplemented with 10% fetal calf serum and 1% pen/strep) + 2.5% BME + pamidronate (1:2000, Merck Calbiochem 506600-10MG) + Annexin CF555 (Biotium 30105A-100µL)) were added to each well. A small z-stack (50 µm) of each gel was acquired over the course of 15-20 h. Imaging data was processed in Imaris.

### Printing of a human breast model, tumor cell seeding and T cell treatment

MCF10A were trypsinized using TryplE for 15 min at 37 °C and collected in F+++, spun down at 300 g and pellets were resuspended in BME. Single MCF10A cells were allowed to form spheroids for two weeks. For printing experiments, two fully confluent plates of MCF10A spheroids were collected as whole spheroids in F+++, spun down and washed in PBS + 0.1 % BSA. Subsequently, spheroids were resuspended in 12 mL 6 % w/v 60 DoM hybrid gel. 2 mL spheroid suspension per vial were added into 20 mm glass printing vials. The cell suspension was allowed to solidify on ice for 20 minutes, while swivelling occasionally to prevent spheroids from sedimentation.

A breast model containing a pocket (Fig. 4D), which is accessible through the nipple, and filled with a gyroid pattern was volumetrically printed at light doses of 160-200 mJ/cm^2. Right after printing, breast models where washed with warm PBS to remove uncured gel. Per breast, 1 well (12WP) of breast cancer organoids (line 13T) was collected in F+++, spun down at 300 g and resuspended in 5 µL 60 DoM hybrid gelMA and injected through the nipple into the pocket. Bioprinted breasts were post-cured for 2 minutes in the UV oven and cultured in 12 well multiwell plates with BCM, which was refreshed twice a week. For T cell experiments, a PDMS mold (Extended Fig. 8B) was used to flip the breast models upside down to allow access to the gyroid pattern. 1 million TEGs were added per print. Flow through was re-added to the top (bottom of the breast) twice a day for three days.

Breast models were fixed in 4% PFA for 1.5 h at 4°C and subsequently washed and stored in PBT. Breast samples were stained for mouse anti-CD3 (BioLegend 300402; 1:500) and rabbit anti-cleaved-caspase-3 (Cell Signaling 9664T; 1:200) in WB2 overnight. The next day, samples were washed 3x 1.5 h with WB2 and subsequently stained with secondary antibody solution including DAPI (1:1000), Phalloidin-Atto700 (1:200), donkey-anti-mouse-AF647 (1:200), donkey-anti-rabbit-AF647 (ThermoFisher Scientific A31573; 1:200) over night. The next day, samples were washed 3x 1.5 h in WB2. Subsequently, breasts were submerged in 4 % low melting agarose (UltraPure LMP Agarose, Invitrogen, 16520-050). After solidification of the agarose, breasts were cut at the vibratome (Leica VT1200S, 0.8 mm/s, 1 mm amplitude) into 350 µm thick sections. In total six boobs were printed containing two different BCO lines. From both donors two breasts were exposed to TEGs. Per breast, two sections were imaged and analysed.

In imaris, segmentation was performed on the breast volume, the tumor volume, MCF10A spheroids, BCOs and T cells. T cell densities within the different compartments were calculated in RStudio.

### YAP staining, imaging & analysis

Gel discs were fixed for 1 h in 4% PFA at 4 °C and then washed and stored in PBT. Samples were embedded in 4% low melting agarose and vibratome cut into cross sections (Extended Fig. 8C) of 350 µm thickness. Per donor three samples per condition (Covalent 100 DoM, hybrid and covalent 60 DoM) were cut. Per sample three regions were stained for YAP (Santa Cruz Biotechnology YAP Antibody (63.7): sc-101199;1:100) in WB2 at 4°C over night. The next day samples were washed 3x 1.5 h in WB2 and then stained for DAPI (1:1000), Phalloidin-AF555 (1:200) and donkey-anti-mouse-AF647 (1:200). Z-stacks of 150-200 µm were acquired at 25 x magnification and regions were chosen that do not include the edges of the samples. In imaris, cell bodies were segmented based on the Phalloidin signal and nuclei were segmented based on the DAPI signal. 3D coordinates of cell bodies and nuclei, as well as YAP mean intensity and cell sphericity were exported to csv files and imported into RStudio. In RStudio based on the x,y,z-coordinates of cell bodies and nuclei, each cell body was assigned one nucleus. Subsequently, YAP intensity in nuclei and cell bodies was analysed and the nuclear/cytoplasmic YAP ratio was calculated per single cell and plotted in relation to sphericity.

### Statistical analysis

Results were displayed as mean ± standard deviation (SD) unless indicated otherwise. Statistical analysis was performed using GraphPad Prism 9 (GraphPad Software, USA). Comparisons between experimental groups was assessed via One-way ANOVA followed by Tukey’s multiple comparisons test (> 2 conditions) or unpaired parametric t test (2 conditions).

Results were considered significant with a p-value < 0.05 and displayed in the graphs as follows: NS = not significant; * = p < 0.05; ** = p < 0.01; *** = p < 0.001; **** = p < 0.0001.

**Extended Figure 8.**
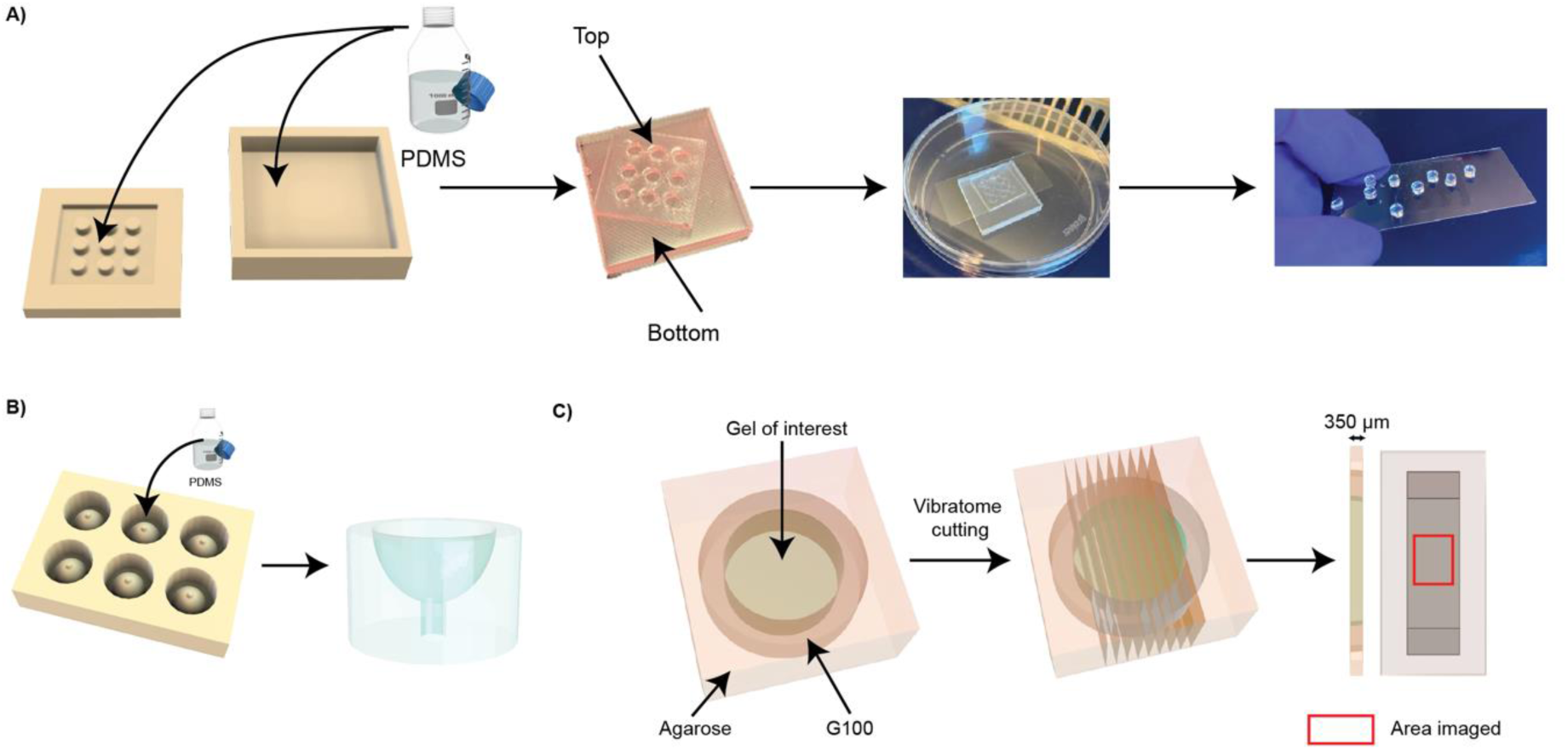
a) Casting approach to make organoid hydrogel discs. b) Casting a PDMS holder to culture bioprinted breasts upside down. c) Cutting and imaging approach to process hydrogel samples stained for YAP.

